# NON-NUCLEOSIDE REVERSE TRANSCRIPTASE INHIBITORS BLOCK THE INCORPORATION OF VIRAL GLYCOPROTEINS INTO HIV-1 VIRIONS

**DOI:** 10.64898/2026.11.10.765689

**Authors:** Abdul A. Waheed, Hanan Ibrahim, Gokul Raghunath, Roshna Vijayan, Gregory B. Melikyan, Eric O. Freed

## Abstract

Non-nucleoside reverse transcriptase inhibitors (NNRTIs) are a key component of antiretroviral therapy that block reverse transcription in newly infected cells. NNRTIs have also been reported to induce premature activation of the HIV-1 protease in virus-producing cells, an activity that has been optimized in a class of NNRTIs known as targeted activators of cell kill (TACK). In this study, we used the TACK compound Pyr01 to investigate how NNRTI-induced premature protease activation affects virion composition and infectivity. We found that Pyr01 treatment markedly reduces the incorporation of the HIV-1 envelope (Env) glycoprotein into virus particles in an Env cytoplasmic tail-independent manner. Pyr01 treatment also disrupts the virion incorporation of foreign viral glycoproteins and causes a substantial loss of the viral enzymes from virions. Virions produced from NNRTI-treated cells exhibit reduced membrane order. The defect in viral glycoprotein incorporation requires an active protease, implicating premature protease-mediated disruption of the Gag lattice and resulting changes in membrane properties as the primary mechanism underlying these effects. These findings reveal that NNRTI-induced premature protease activation during virus assembly fundamentally changes the protein composition and membrane order of HIV-1 virions to markedly reduce particle infectivity. These results have implications for the use of TACK compounds therapeutically and provide basic insights into the properties of the viral membrane that promote viral glycoprotein incorporation.

## Introduction

The HIV-1 Gag and GagPol polyprotein precursors provide the structural components and viral enzymes required for virus particle assembly, budding, and maturation. The Gag polyprotein precursor, Pr55Gag, contains several functional domains: the matrix (MA) domain targets Gag to the plasma membrane and facilitates envelope glycoprotein (Env) incorporation into the virion; the capsid (CA) domain drives Gag multimerization to form the immature hexameric Gag lattice; the nucleocapsid (NC) domain selectively binds and packages the viral genomic RNA and also promotes particle assembly; the p6 domain recruits the host endosomal sorting complex required for transport (ESCRT) machinery that drives membrane scission at the neck of the budding particle to allow virion release; and the spacer peptides 1 and 2 (SP1 and SP2) contribute to regulated Gag assembly, processing, and maturation (see reviews^1–4^). The GagPol precursor, Pr160GagPol, is generated by a programmed -1 ribosomal frameshift in approximately 5% of translation events and contains the domains for the viral enzymes protease (PR), reverse transcriptase (RT), and integrase (IN). All three viral enzymes are oligomeric in their active state. PR functions as an obligate homo-dimer, with the active site formed by the dimer interface^5–8^. Gag and GagPol co-assemble at the inner leaflet of the plasma membrane to form immature virions. During the late stages of assembly and budding, GagPol interactions promote PR dimerization and activation. Activated PR then cleaves Gag and GagPol polyproteins in a highly orchestrated, sequential cascade, initiating the transition from a non-infectious immature particle to an infectious mature virion^1,3,9^.

The spatio-temporal timing of PR activation is a critical determinant of HIV-1 replication; either delaying PR activation or inducing premature, intracellular PR activation disrupts the timing of Gag and GagPol processing, leading to defective virion assembly, aberrant maturation, and a loss of infectivity^10^. Early studies demonstrated that the activity of HIV-1 PR is initiated at the host cell membrane before virus release^11^. More recently, a variety of imaging approaches were used to confirm that HIV-1 PR is activated within producer cells during assembly and budding prior to particle release^12^.

HIV-1 assembly takes place in highly ordered, lipid raft-like plasma membrane microdomains enriched in cholesterol and saturated lipid species^13,14^; for reviews, see^15,16^. The phospholipid PI(4,5)P_2_ plays a key role in trafficking of Gag to the inner leaflet of the host cell plasma membrane^17–19^ via direct MA-PI(4,5)P_2_ interactions^20^. PI(4,5)P_2_ has also been shown to be required for the stable retention of assembled Gag complexes at the plasma membrane^21^. Consistent with a role for PI(4,5)P_2_, cholesterol, and saturated lipids in HIV-1 assembly, the lipid composition of viral membrane is distinct from that of the bulk plasma membrane of producer cells and is enriched in cholesterol, saturated lipid species, and PI(4,5)P ^22–28^. Several studies provided evidence that HIV-1 Gag actively reorganizes the plasma membrane at sites of assembly to generate a specialized lipid nanodomain^29,30^. Using model membranes, it was demonstrated that Gag assembly promotes PI(4,5)P□ clustering and recruits cholesterol into these domains^30^ and that Gag assembly reduces the mobility of PI(4,5)P□ and cholesterol at assembly sites^29^. Gag oligomerization induces changes in membrane organization at assembly sites that drive lipid-phase-dependent partitioning of membrane proteins^31^. These findings support a model in which Gag multimerization drives the formation of distinct PI(4,5)P□ -and cholesterol-enriched membrane microdomains that serve as platforms for HIV-1 assembly. Thus, Gag functions not only as the structural scaffold for particle assembly but also organizes the local membrane environment, contributing to the selective lipid and protein composition of budding HIV-1 particles.

HIV-1 Env is synthesized as a precursor polyprotein, gp160, that is cleaved by host furin-like proteases during trafficking through the Golgi apparatus into the non-covalently associated gp120 and gp41 subunits that make up the mature, hetero-trimeric Env complex. The gp41 subunit of HIV-1 and other primate lentiviruses has a cytoplasmic tail (CT) that is very long (∼150 amino acids) compared with the CTs of most other viral glycoproteins. The mechanism by which Env is incorporated into HIV-1 virions remains incompletely understood but it is well established that the long gp41 CT is required in most cell types for Env incorporation and the CT thought to be engaged by the MA component of the immature Gag lattice during virus assembly^32–46^. However, foreign viral glycoproteins – e.g., vesicular stomatitis virus G (VSV-G), murine leukemia virus (MLV) Env, SARS-CoV-2 spike (S), and others – can be incorporated into HIV-1 particles in a process known as pseudotyping^47^. These foreign viral glycoproteins have relatively short CTs that are not thought to directly engage the immature HIV-1 Gag lattice but rather are incorporated non-specifically during the acquisition of host cell plasma membrane by the virus during assembly. In addition to this non-specific mechanism of incorporation, in many cases these foreign viral glycoproteins are also postulated to be enriched at sites of assembly because of the unique lipid environment (described above) created by the assembling Gag lattice^29,30^.

Mature RT is an asymmetric p66/p51 heterodimer; the p66 subunit contains both the active polymerase and RNase H domains, whereas p51 is generated by proteolytic removal of the RNase H domain from p66 and primarily provides structural support to the heterodimeric enzyme. Non-nucleoside RT inhibitors (NNRTIs) block HIV-1 RT by binding to an allosteric hydrophobic pocket in the p66 subunit of the enzyme, inducing conformational changes that impair its DNA polymerase activity^48^. Certain NNRTIs, particularly efavirenz (EFV), enhance RT dimerization^49^. Subsequent work confirmed that EFV promotes RT homodimerization and accelerates the proteolytic processing of a model Pol precursor^50^. These findings were extended to demonstrate that potent NNRTIs enhance GagPol interactions, leading to premature PR activation and aberrant pre-budding polyprotein cleavage that results in defective virion assembly and reduced virus production^51^. Thus, in addition to inhibiting RT polymerase activity post-infection, NNRTIs can disrupt the late-stages of HIV-1 replication by promoting GagPol oligomerization and prematurely activating PR. NNRTI-induced premature PR activation can selectively kill HIV-1 expressing CD4+ T cells^52,53^ by triggering CARD8 inflammasome activation, leading to caspase-1-dependent pyroptosis and selective killing of HIV-1-infected cells^54^. This cell killing activity has been optimized in a class of compounds referred to as targeted activators of cell kill (TACK)^55^. High-resolution structural studies have provided insights into the molecular basis for how NNRTIs stabilize RT-mediated GagPol interactions, thereby promoting premature PR activation and potentiating CARD8 inflammasome activation^56^.

In this study, we investigated the consequences of NNRTI-induced premature HIV-1 PR activation on virion assembly, maturation and infectivity. We observed a marked reduction in the infectivity of HIV-1 particles produced from NNRTI-treated cells, which correlated with a severe reduction in Env incorporation into virions. Importantly, the Env incorporation defect was rescued by PR inactivation or expression of cleavage-resistant Gag, implicating premature Gag cleavage and consequent disruption of the Gag lattice as the underlying cause. NNRTI treatment of virus-producing cells also caused a loss of the Pol-derived enzymes (PR, RT, and IN) from released particles. Together, these findings identify previously uncharacterized consequences of NNRTI-induced premature PR activation on HIV-1 particle composition and infectivity.

## Results

### Treatment of virus-producing cells with NNRTIs impairs infectivity of HIV-1 virions

Certain NNRTIs, including Pyr01 and efavirenz (EFV), have been shown to enhance GagPol dimerization in HIV-1-producing cells, resulting in premature activation of PR during virus assembly^49–51,55^. To investigate the consequences of NNRTI treatment on the late stages of the HIV-1 replication cycle, virus-producing 293T cells were treated with increasing concentrations of Pyr01, ranging from 0.1 to 0.4 μM. Virus-containing supernatants were collected, and virus production and infectivity were assessed. Western blot analysis of cell lysates showed that increasing concentrations of Pyr01 enhance Gag processing in producer cells, as evidenced by a decrease in Pr55Gag levels and increased levels of the Gag processing intermediate p41 and mature capsid (CA) (Fig. 1A, upper panel). These findings are consistent with Pyr01-induced GagPol dimerization, leading to premature activation of HIV-1 PR and enhanced Gag processing. Notably, at these concentrations, Pyr01 had no measurable effect on virus particle production (Fig. 1A, lower panel).

**Fig. 1.**
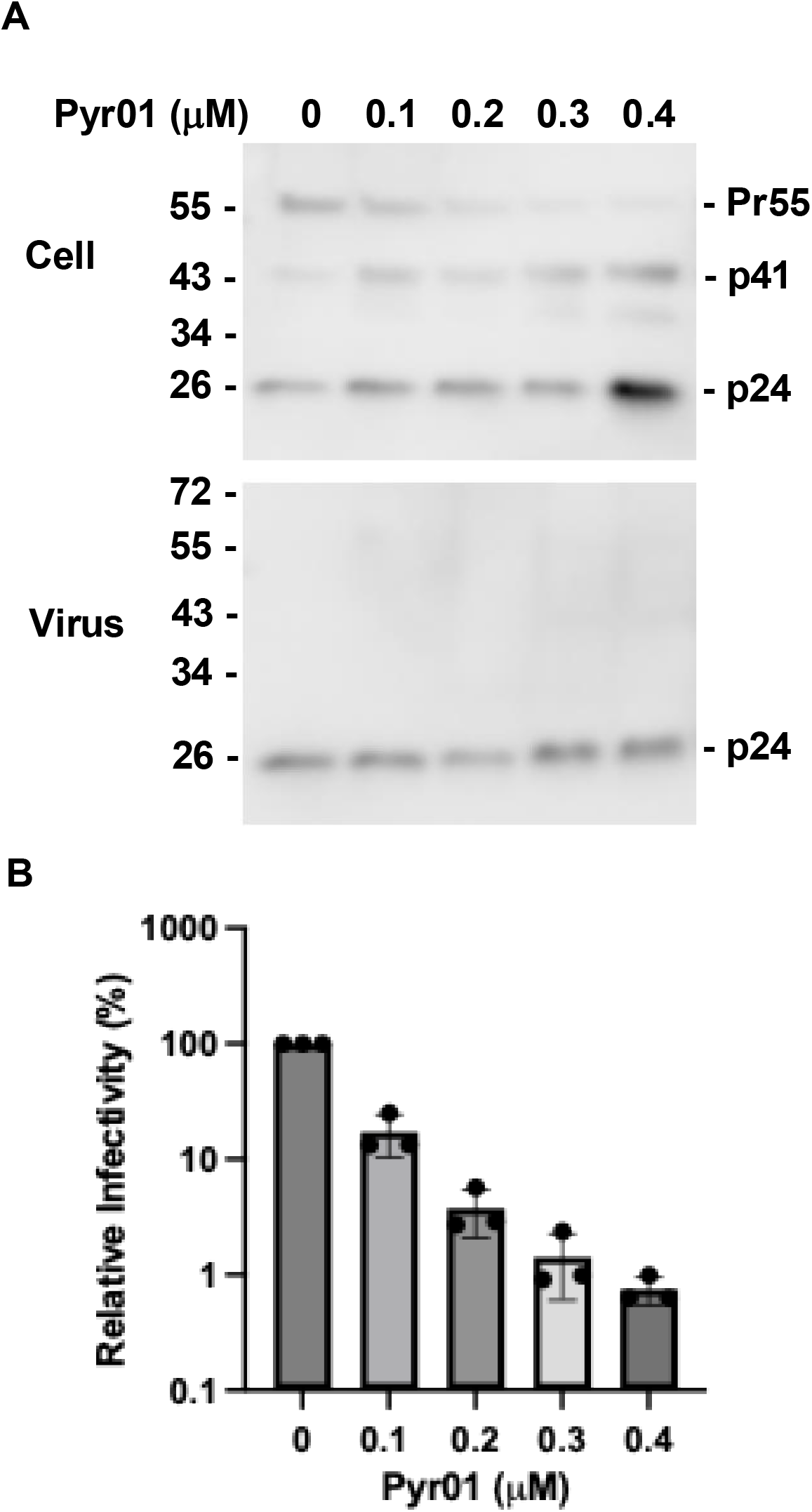
Treatment of virus-producing cells with Pyr01 impairs infectivity of HIV-1 virions. **(A).** 293T cells were transfected with the HIV-1 proviral clone pNL4-3, and 6-8 h posttransfection, treated with Pyr01 at 0, 0.1, 0.2, 0.3 and 0.4 μM. After 16-20 h of treatment, cell and virus lysates were prepared and analyzed by western blotting using HIV-Ig to detect the Gag precursor Pr55Gag (Pr55), Gag processing intermediates (p49 and p41), and mature CA (p24). The positions of molecular mass standards (in kDa) are indicated on the left of each blot. **(B)**. TZM-bl cells were infected with serial dilutions of virus-containing supernatants prepared as in panel A. Luciferase activity was measured 48 h postinfection, and infectivity of virus produced in the absence of Pyr01 was normalized to 100%. Data shown are the mean ± SD from three independent experiments.

We next examined the infectivity of virus produced from Pyr01-treated cells using the TZM-bl indicator cell line. Strikingly, Pyr01 treatment caused a severe, dose-dependent reduction in viral infectivity, with a more than 100-fold decrease at the highest concentration tested (Fig. 1B). Because NNRTIs inhibit HIV-1 replication by blocking reverse transcriptase (RT) activity during infection, we asked whether residual Pyr01 carried over in the virus-containing supernatant could account for the observed loss of infectivity. To address this possibility, Pyr01 was added directly to TZM-bl cells at concentrations equivalent to the estimated final concentrations present during infection with virus produced from Pyr01-treated cells. Under these conditions, Pyr01 reduced infectivity by less < 2-fold, even at the highest final concentration tested (11 nM), which is well below its reported RT inhibitory IC_50_ of approximately 40 nM, and is consistent with its expected antiviral activity (Fig. S1). These results indicate that the marked loss of infectivity of virus produced in the presence of Pyr01 (Fig. 1) cannot be explained solely by inhibition of reverse transcription during target cell infection.

To determine whether the defect in NNRTI treatment of virus-producer cells is specific for HIV-1 Env-mediated entry, we examined the infectivity of VSV-G-pseudotyped HIV-1 produced in the presence of Pyr01. Similar to wild-type (WT) HIV-1, the infectivity of VSV-G-pseudotyped HIV-1 was reduced by more than 100-fold when virus-producing cells were treated with 0.3 and 0.4 μM Pyr01 (Fig. S2). Collectively, these findings demonstrate that Pyr01 treatment of virus-producing cells at the concentrations tested markedly impairs the infectivity of WT and VSV-G-pseudotyped HIV-1 without affecting virus production.

### NNRTs severely reduce the incorporation of viral envelope glycoproteins

To investigate the mechanism(s) responsible for the reduction in virus particle infectivity resulting from treatment of virus-producing cells with Pyr01, we examined the content of HIV-1 virions by western blot analysis. Virus-producing 293T cells were treated with 0.25 or 0.5 μM Pyr01, and cell and virus lysates were analyzed by western blotting using antibodies that detect HIV-1 Gag and Env. Pyr01 treatment did not alter the cellular expression of HIV-1 Env (Fig. 2A, upper left panel); however, it markedly reduced the incorporation of Env into HIV-1 virions (Fig. 2A, lower left panel). Consistent with the results shown in Fig. 1, virus production was not affected at these concentrations of Pyr01 (Fig. 2A, middle left panel). To determine whether the loss of Env incorporation is specific to Pyr01 or represents a more general consequence of NNRTI-induced premature PR activation, we examined the effects of EFV, another NNRTI that promotes GagPol dimerization. Virus-producing cells were treated with 10 or 20 μM EFV, and cell and viral lysates were analyzed by western blotting. Similar to Pyr01, EFV did not affect cellular Gag or Env expression but markedly decreased Env incorporation into HIV-1 virions (Fig. 2A, lower right panel). Env incorporation into virions was quantified by measuring the ratio of virion-associated gp41/p24 (CA). This analysis demonstrated an approximately 30-fold reduction in Env incorporation following treatment of virus-producing cells with 0.5 μM Pyr01 and an approximately 25-fold reduction following treatment with 20 μM EFV (Fig. 2B).

**Fig. 2.**
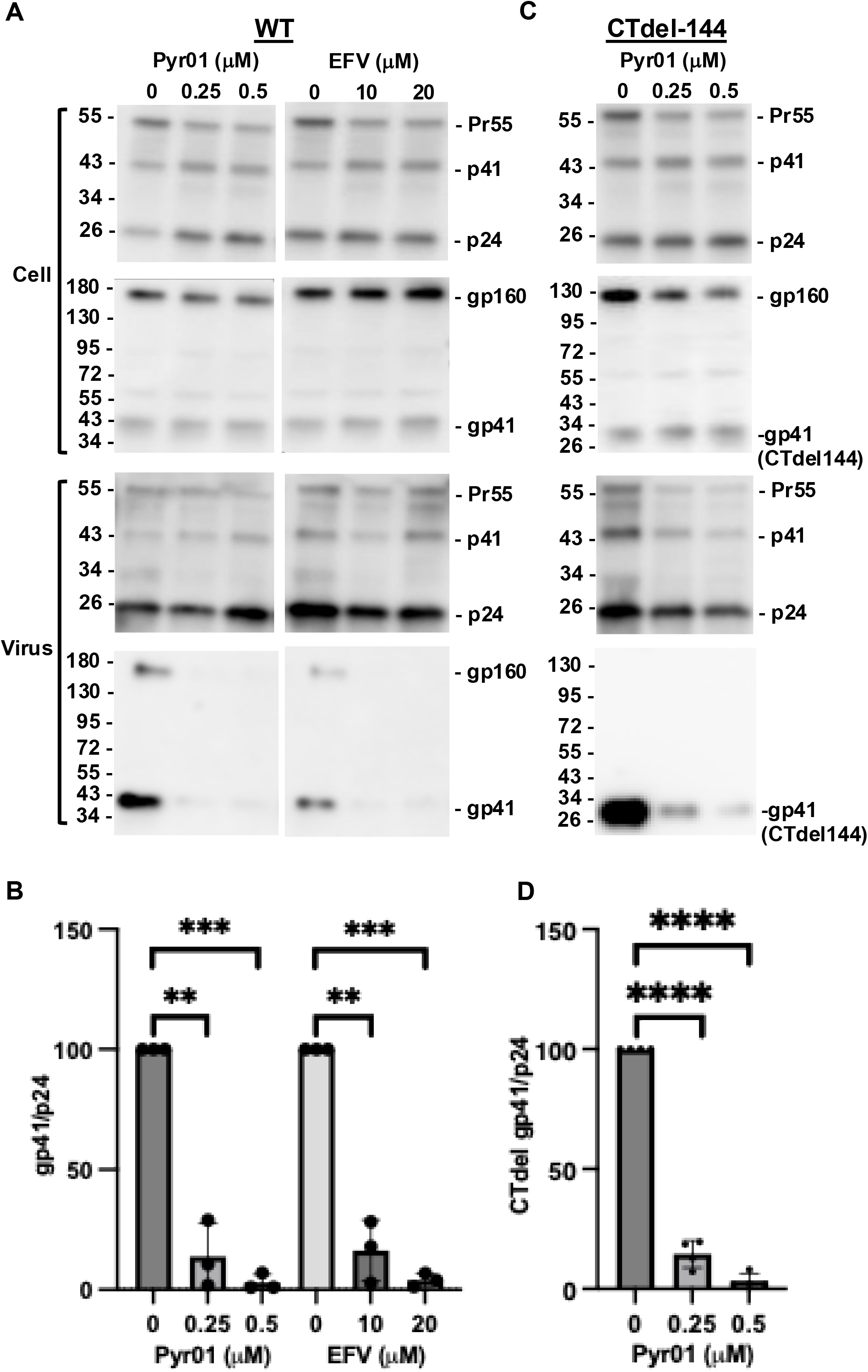
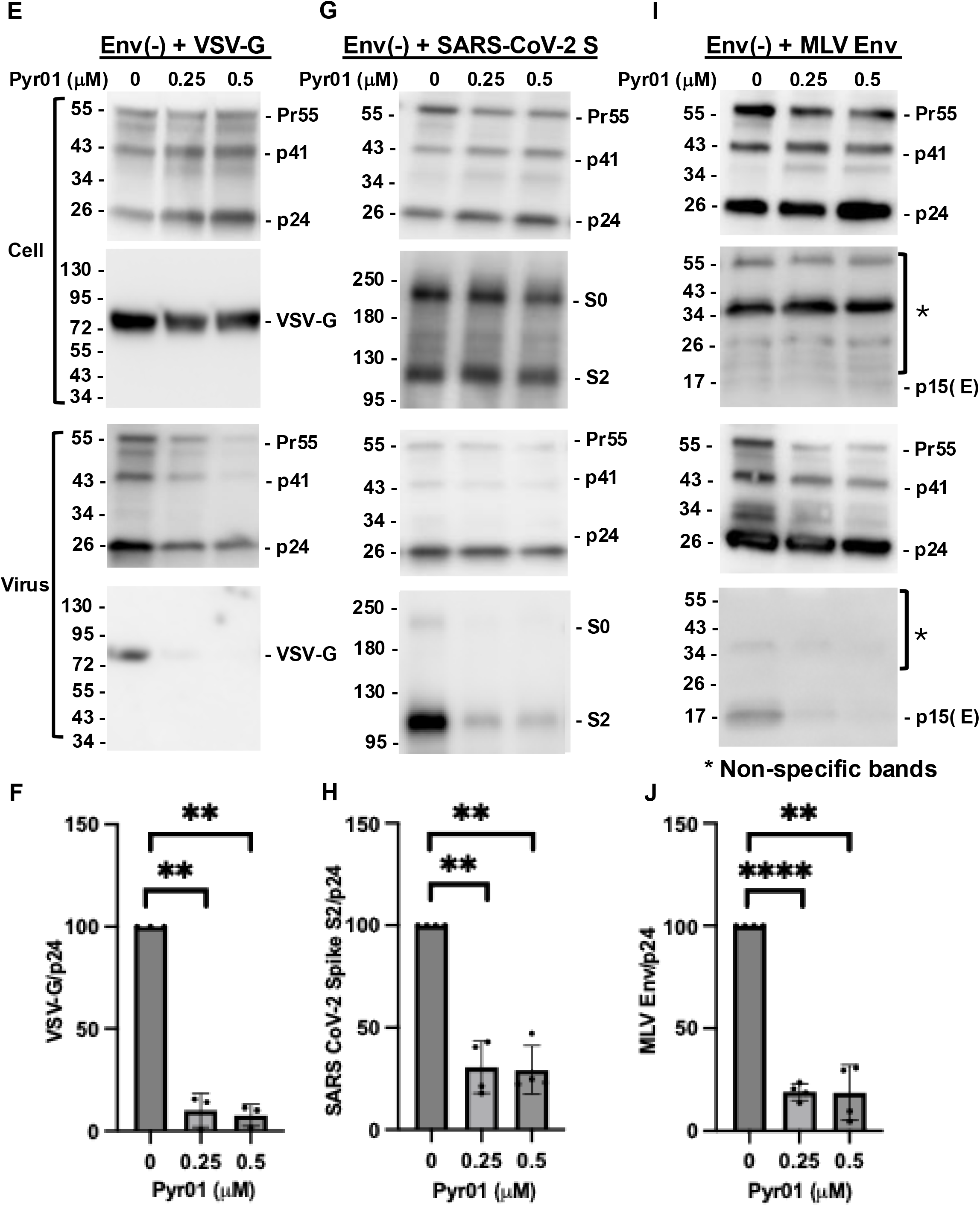
NNRTs severely reduce the incorporation of viral envelope glycoproteins. 293T cells were transfected with WT pNL4-3 **(A, B)** or the gp41 cytoplasmic-tail truncation mutant CTdel-144 **(C, D)** or co-transfected with the Env-defective molecular clone pNL4-3/KFS [Env (-)] and expression vectors encoding VSV-G **(E, F)**, or SARS-CoV-2 spike **(G, H)**, or MLV Env **(I, J)**. Six to eight h posttransfection, cells were treated with Pyr01 at 0, 0.25 and 0.5 μM or with efavirenz (EFV) at 0, 10 and 20 μM. After 16-20 h of treatment, culture supernatants were collected, and cell and virus lysates were analyzed by western blotting using HIV-Ig as described in Fig. 1A. Envelope glycoproteins were detected using the following antibodies: anti-HIV-1 gp41 monoclonal antibody 2F5 **(A and C)**, anti-VSV-G **(E)**, anti-SARS-CoV-2 spike **(G)**, and anti-MLV Env **(I)**. The positions of molecular mass standards (in kDa) are indicated on the left of each blot. Virion-associated envelope glycoprotein and p24 levels were quantified using Image Lab software version 6.0.1 (Bio-Rad). The ratio of envelope glycoprotein to p24 (CA) protein was calculated with untreated set to 100% **(B, D, F, H, J)**. Data shown are the mean ± SD from three **(B, F)** or four **(D, H, J)** independent experiments. *P < 0.05, **P < 0.01, ***P < 0.001, ****P < 0.0001; unpaired t-test.

Next, we asked whether the reduction in HIV-1 Env incorporation induced by NNRTI treatment of virus-producing cells is specific to HIV-1 Env or represents a broader defect affecting the incorporation of heterologous viral glycoproteins. We also sought to determine whether truncation of the gp41 cytoplasmic tail affects the NNRTI-induced defect in HIV-1 Env incorporation. To address these questions, we measured the incorporation of several viral envelope glycoproteins into HIV-1 particles – VSV-G, SARS-CoV-2 spike, and amphotropic murine leukemia virus (MLV) Env – and tested a cytoplasmic tail-truncated HIV-1 Env mutant (CTdel-144;^38^ (Fig. S3). Unlike WT HIV-1 Env, which possesses an unusually long cytoplasmic tail of approximately 150 amino acids, these glycoproteins contain substantially shorter cytoplasmic tails: CTdel-144 retains only ∼ six cytoplasmic tail residues, whereas VSV-G, SARS-CoV-2 spike, and amphotropic MLV Env contain cytoplasmic tails of 21, 39, and 32 amino acids, respectively (Fig. S3). Western blot analysis of virions demonstrated that Pyr01 treatment of virus-producing cells markedly reduces the incorporation of CTdel-144 HIV-1 Env, with an approximately 30-fold decrease in the gp41/p24 (CA) ratio at 0.5 μM Pyr01 (Fig. 2C-D). Similarly, incorporation of VSV-G was reduced by approximately 14-fold following treatment with either Pyr01 or EFV (Fig. 2E-F and Fig. S4). Incorporation of the SARS-CoV-2 spike protein was also significantly reduced, although to a lesser extent (∼ three-fold), following treatment with 0.5 μM Pyr01 (Fig. 2G-H). The incorporation of amphotropic MLV Env was also affected by Pyr01 treatment (Fig. 2I-J). It is noteworthy that neither Pyr01 nor EFV altered the cellular expression levels of CTdel-144 HIV-1 Env, VSV-G, MLV Env or SARS-CoV-2 spike. These findings demonstrate that the defect in envelope glycoprotein incorporation induced by NNRTIs is not restricted to HIV-1 Env and cannot be attributed solely to the unusually long cytoplasmic tail of HIV-1 Env. Instead, NNRTI treatment of virus-producing cells impairs the incorporation of multiple viral envelope glycoproteins, including HIV-1 Env, CTdel-144 HIV-1 Env, VSV-G, MLV Env and SARS-CoV-2 spike. These findings suggest that premature HIV-1 PR activation may perturb one or more mechanisms governing envelope glycoprotein incorporation into budding HIV-1 particles.

### The NNRTI-mediated reduction in envelope glycoprotein incorporation requires an active PR

Because NNRTIs promote GagPol dimerization in virus-producing cells, resulting in premature HIV-1 PR activation, we hypothesized that inhibition of PR activity might rescue the defect in Env incorporation. To test this, virus-producing 293T cells were treated with Pyr01 in the absence or presence of the HIV-1 PR inhibitor saquinavir (SQV). As shown in Fig. 3A-B, inhibition of PR activity by SQV completely restored the incorporation of HIV-1 Env into virions (compare virus-associated gp41 levels in lanes 2 and 3 with lanes 5 and 6), indicating that the Pyr01-induced defect in Env incorporation depends on PR activity. We next examined whether SQV could similarly rescue the incorporation defects observed with truncated HIV-1 Env and with a heterologous viral envelope glycoproteins. Consistent with the results obtained for WT HIV-1 Env, Pyr01 treatment did not significantly reduce the incorporation of CTdel-144 Env into virions in the presence of SQV (Fig. S5). Likewise, the decrease in VSV-G incorporation caused by Pyr01 was significantly rescued by SQV treatment (Fig. 3C-D). These results demonstrate that inhibition of PR activity reverses the Pyr01-mediated incorporation defects of multiple viral envelope glycoproteins.

**Fig. 3.**
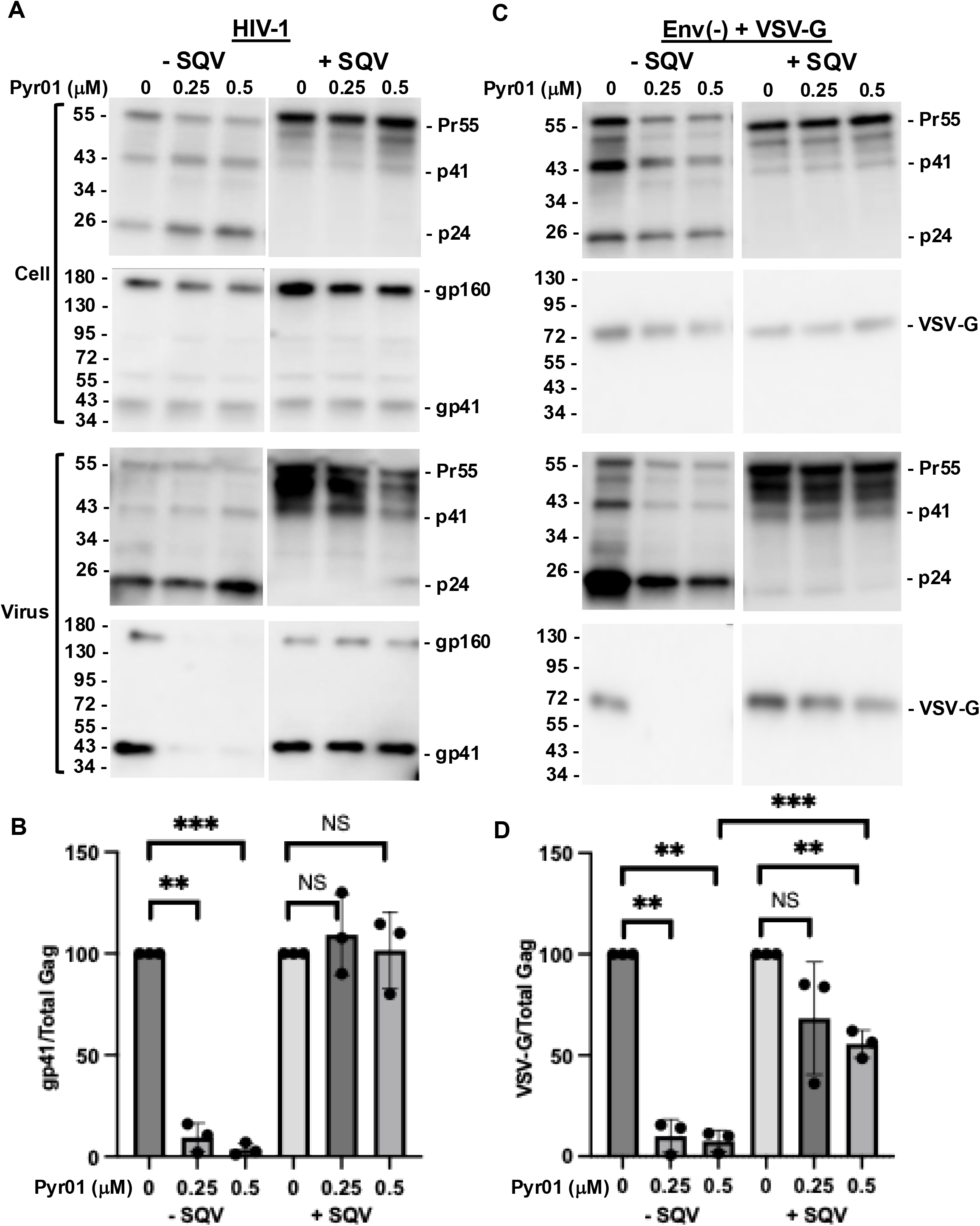
Inhibition of PR activity with SQV reverses the Pyr01-mediated envelope glycoprotein incorporation defect. 293T cells were transfected with WT pNL4-3 **(A,B)** or co-transfected with the Env-defective molecular clone pNL4-3/KFS [Env (-)] and VSV-G expression vector **(C, D)**. Six to eight h posttransfection, cells were treated with Pyr01 at 0, 0.25 or 0.5 μM in the presence or absence of 0.5 μM of the protease inhibitor saquinavir (SQV). After 16-20 h of treatment, culture supernatants were collected, and cell and virus lysates were analyzed by western blotting using HIV-Ig, and anti-gp41 monoclonal antibody 2F5 or anti-VSV-G antibody, as described in Fig. 2. The positions of molecular mass standards (in kDa) are indicated on the left of each blot. **(B,D)** Virion-associated gp41 and VSV-G and p24 levels were quantified and the ratio of envelope glycoprotein to p24 (CA) protein was calculated with untreated set to 100%, as described in the Fig. 2 legend. Data shown are the mean ± SD from three independent experiments. **P < 0.01, ***P < 0.001; unpaired t-test.

To independently confirm the requirement for PR activity in the ability of Pyr01 to reduce envelope glycoprotein incorporation, we tested whether genetic inactivation of PR produced a similar phenotype as SQV in reversing the ability of Pyr01 to reduce envelope glycoprotein incorporation. A derivative of NL4-3 carrying a catalytic-site mutation in PR (D25N;^57^) was analyzed in the presence or absence of Pyr01. As shown in Fig. 4 (middle panel), Pyr01 did not impair HIV-1 Env incorporation in the context of the catalytically inactive PR. In addition, we examined an HIV-1 mutant lacking GagPol expression (404-bp deletion within Pol;^58^). Consistent with the result obtained with the clone containing an inactive PR, Pyr01 treatment of virus-producing cells did not reduce Env incorporation into virions (Fig. 4, right panel). Together, these pharmacologic and genetic approaches demonstrate that the Pyr01-induced defect in envelope glycoprotein incorporation requires an active PR, supporting the hypothesis that this effect of Pyr01 is linked to its ability to prematurely activate PR by promoting GagPol dimerization.

**Fig. 4.**
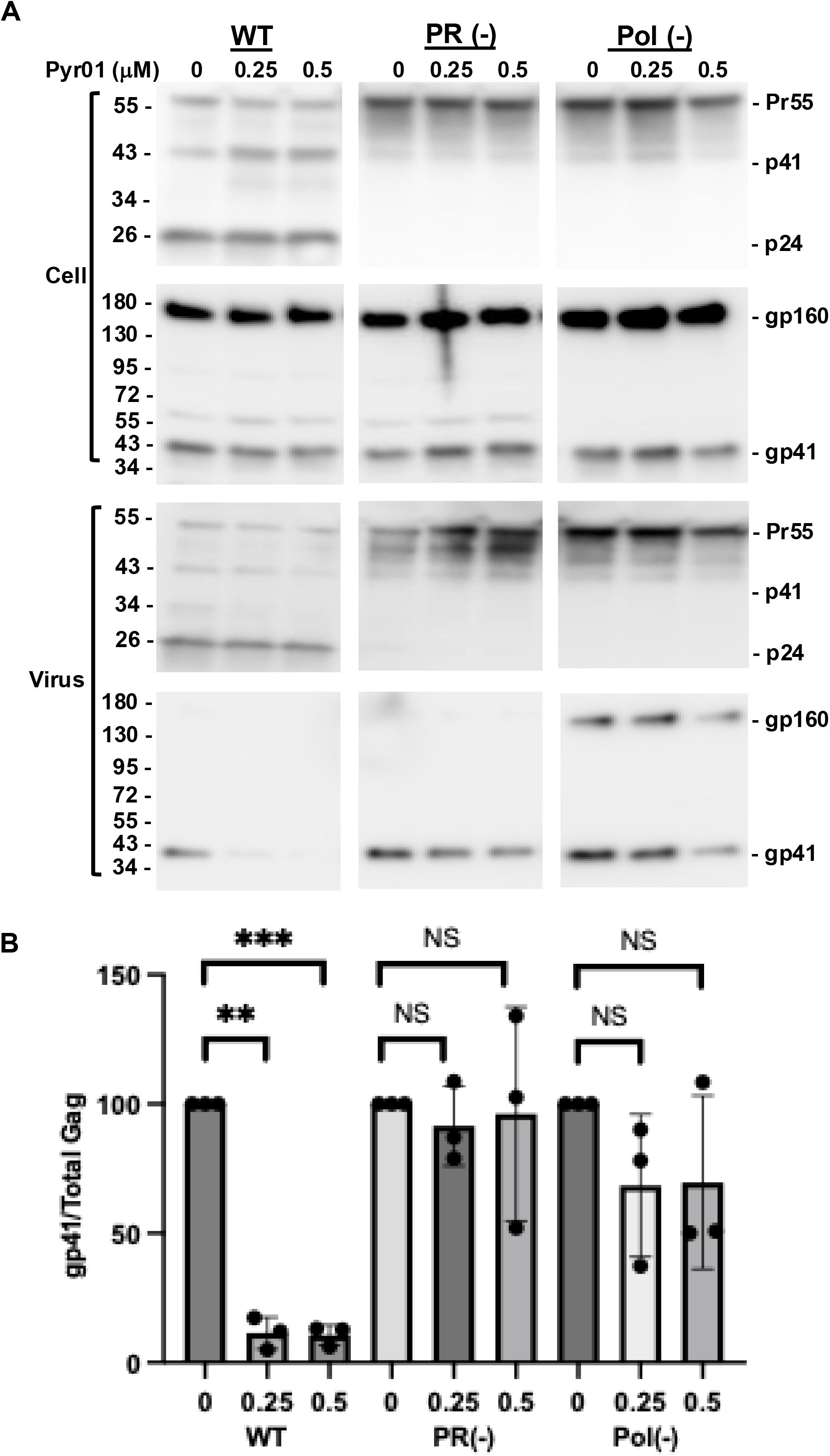
Inactivating or deleting PR prevents Pyr01-mediated reduction in Env incorporation. **(A)** 293T cells were transfected with WT pNL4-3 (WT), a PR-inactive mutant [PR(-)], or a Pol deletion mutant [Pol(-)]. Six to eight h posttransfection, cells were treated with 0, 0.25 and 0.5 μM Pyr01. After 16-20 h of treatment, culture supernatants were collected and cell and virus lysates were analyzed by western blotting using HIV-Ig and anti-gp41 2F5 antibodies, as described in the Fig. 2 legend. The positions of molecular mass standards (in kDa) are indicated on the left of each blot. **(B)** Virion-associated gp41 and p24 levels were quantified and the ratio of gp41 to p24 (CA) protein was calculated with untreated set to 100%, as described in the Fig. 2 legend. Data shown are the mean ± SD from three independent experiments. NS, not significant, **P < 0.01, ***P < 0.001; unpaired t-test.

### HIV-1 Env incorporation into VLPs produced by a Gag mutant in which PR cleavage sites have been mutationally inactivated is not affected by NNRTI treatment

The results presented above demonstrate that the NNRTI-induced envelope glycoprotein incorporation defects depend on PR activity that cleaves Gag during virion maturation. We therefore next examined the effect of mutating Gag cleavage sites on the Pyr01-induced defect in envelope glycoprotein incorporation. To this end, we analyzed a panel of Gag cleavage site mutants – MA/CA, MA/SP1, MA/NC and MA/p6 – in which the cleavage site between the indicated Gag domains were mutated (Fig. S6A;^59^). Importantly, these mutants retain an active PR capable of processing GagPol-derived Pol proteins IN, RT, and PR (Fig. S6B), allowing us to separate the effect of Gag cleavage from that of PR activity itself. Interestingly, the MA/p6 mutant, in which all Gag cleavage sites are mutated, reversed the Pyr01-induced defect in Env incorporation, whereas the partial Gag cleavage site mutants (MA/CA, MA/SP1 and MA/NC) showed little or no recue (Fig. 5, Fig. S7). These findings demonstrate that preventing Gag cleavage is sufficient to restore Env incorporation despite an active PR, indicating that PR activity alone is not responsible for the defect. Instead, the data suggest that premature Gag cleavage, triggered by NNRTI-induced PR activation, disrupts the Gag lattice required for efficient Env incorporation into assembling virions.

**Fig. 5.**
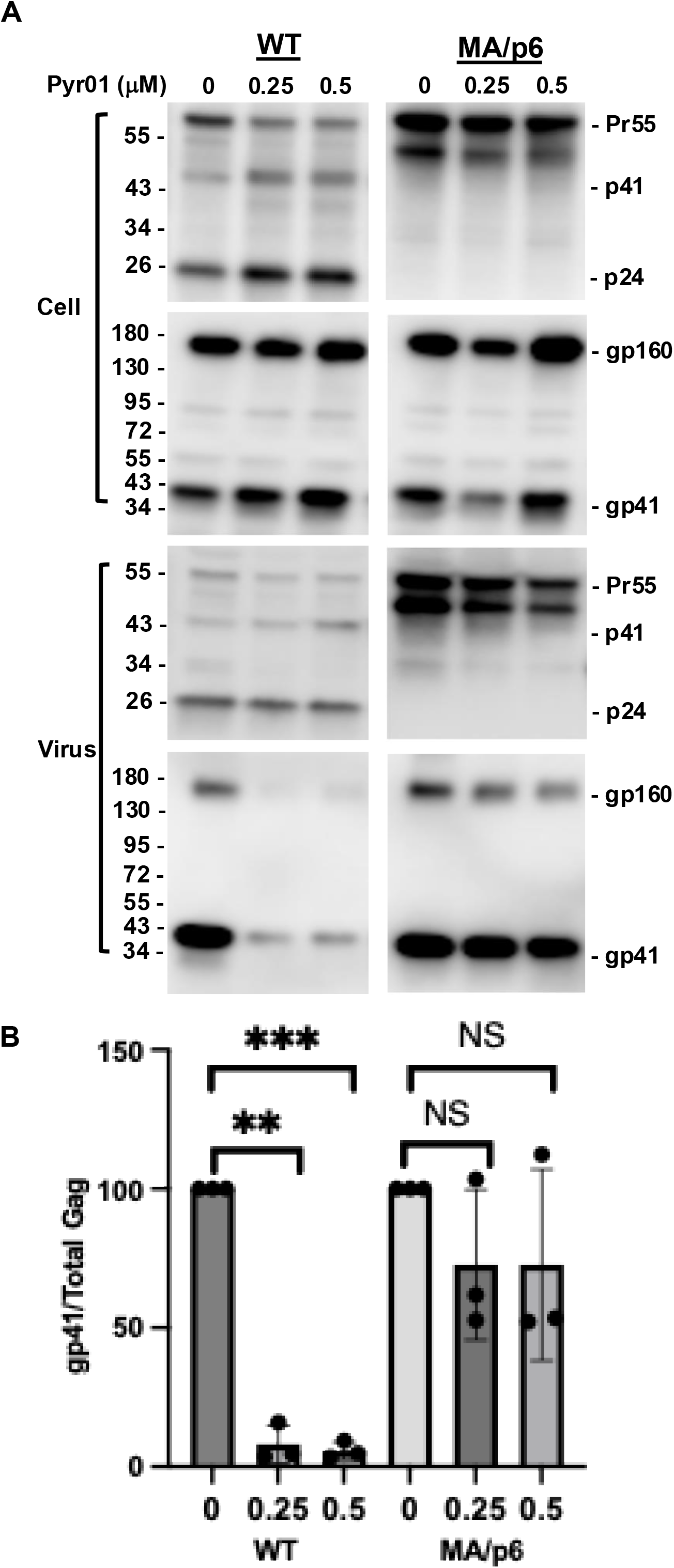
Gag cleavage-site mutant MA/p6 prevents the Pyr01-induced defect in Env incorporation. **(A)** 293T cells were transfected with WT pNL4-3 (WT) or the Gag cleavage-site mutant pNL-MA/p6 (MA/p6). Six to eight h posttransfection, cells were treated with Pyr01 at the indicated concentrations. After 16-20 h of treatment, culture supernatants were collected and cell and virus lysates were analyzed by western blotting using HIV-Ig and anti-gp41 2F5 antibodies, as described in the Fig. 2 legend. The positions of molecular mass standards (in kDa) are indicated on the left of each blot. The positions of molecular mass standards (in kDa) are indicated on the left of each blot. **(B)** Virion-associated gp41 and p24 levels were quantified and the ratio of gp41 to p24 (CA) protein was calculated with untreated set to 100%, as described in the Fig. 2 legend. Data shown are the mean ± SD from three independent experiments. NS -not significant, **P < 0.01, ***P < 0.001; unpaired t-test.

### Pyr01 marginally reduces cell-surface expression of HIV-1 Env and has no effect on VSV-G surface expression

The reduction in viral glycoprotein incorporation into virions induced by NNRTI treatment could potentially be caused by decreased cell-surface expression. To test this possibility, cells were transfected with either WT pNL4-3 or contransfected with the Env(-) derivative pNL4-3/KFS and the VSV-G expression vector and cell-surface expression of HIV-1 Env and VSV-G with and without Pyr01 treatment was measured by flow cytometry. The results indicated that Pyr01 treatment caused a small but statistically significant reduction in the percentage HIV-1 Env-positive cells but had no significant effect on the percentage of VSV-G-positive cells (Fig. S8A). Notably, Pyr01 treatment resulted in only a minimal reduction in the mean fluorescence intensity of either HIV-1 Env or VSV-G (Fig. S8B). These data suggest that the profound effects of NNRTI treatment of virus-producing cells on HIV-1 Env and VSV-G incorporation into virus particles cannot be explained by reduced cell-surface expression.

### Pyr01 induces a defect in Env incorporation across multiple HIV-1 isolates

The experiments described thus far were performed using the laboratory-adapted, CXCR4-tropic HIV-1 molecular clone NL4-3. To determine whether the effect of Pyr01 treatment on Env incorporation is conserved across genetically diverse HIV-1 strains, we extended our analysis to the CCR5-tropic molecular clone NL(AD8) and two subtype C molecular clones, K3016 and ZM247. 293T cells were transfected with these molecular clones in parallel with NL4-3 and treated with Pyr01. Env incorporation into released virions was analyzed by western blotting, and the gp41/p24 ratio was quantified. Consistent with our findings for NL4-3, Pyr01 markedly reduced Env incorporation into virions (Fig. 6A). Quantification of the gp41/p24 ratio revealed a comparable reduction in Env incorporation for NL(AD8), K3016, and ZM247 (Fig. 6B). Together, these results demonstrate that Pyr01 impairs Env incorporation across genetically diverse HIV-1 molecular clones.

**Fig. 6.**
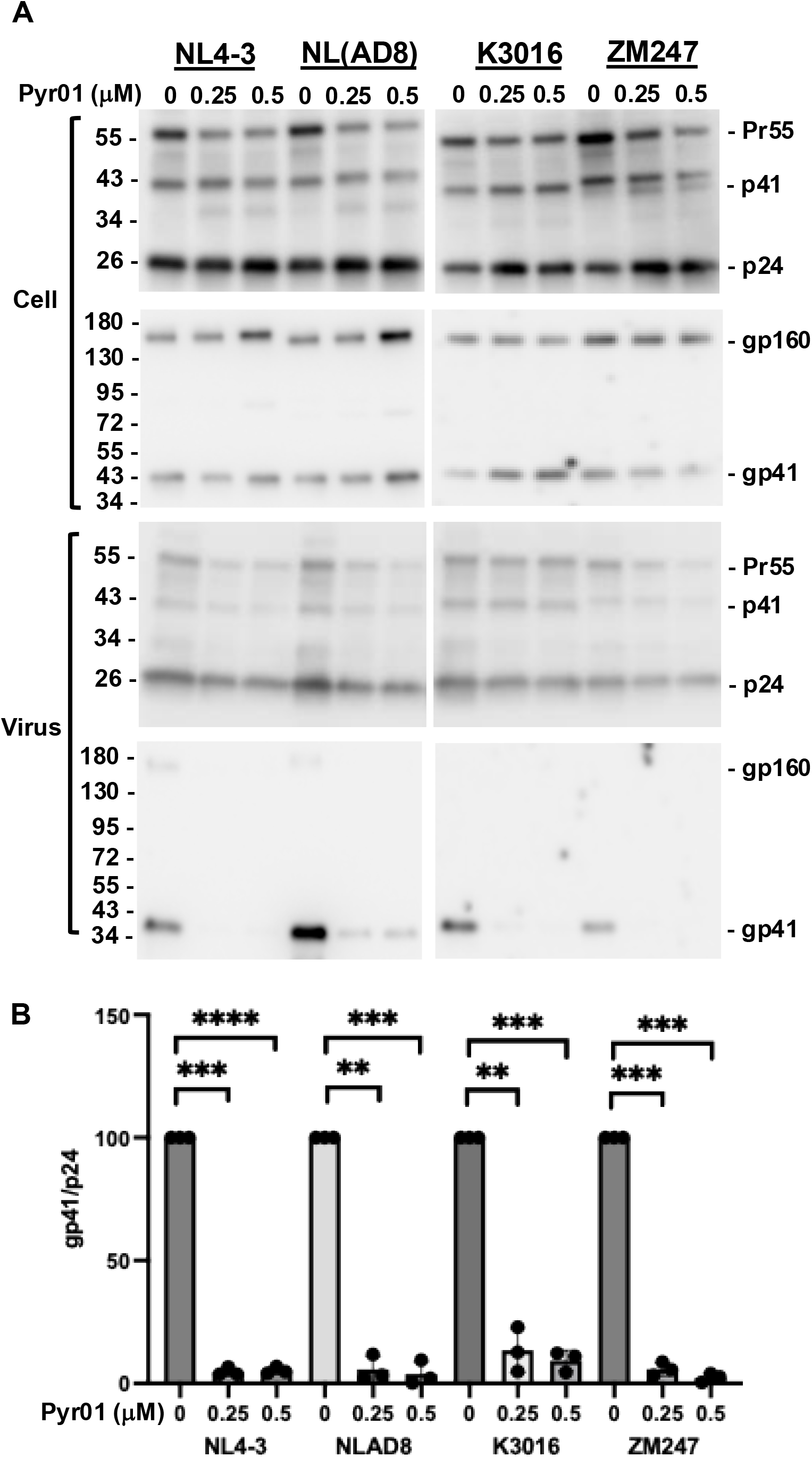
Pyr01 impairs Env incorporation across multiple HIV-1 isolates. **(A)** 293T cells were transfected with pNL4-3, the macrophage-tropic molecular clone pNL(AD8), the HIV-1 subtype C transmitted/founder molecular clone K3016, or the subtype C molecular clone ZM247. Six to eight h posttransfection, cells were treated with Pyr01 at the indicated concentrations. After 16-20 h of treatment, culture supernatants were collected and cell and virus lysates were analyzed by western blotting using HIV-Ig and anti-gp41 2F5 [(pNL4-3 and pNL(AD8)] or anti-gp41 10E8 (K3016 and ZM247) antibodies. The positions of molecular mass standards (in kDa) are indicated on the left of each blot. **(B)** Virion-associated gp41 and p24 levels were quantified and the ratio of gp41 to p24 (CA) protein was calculated with untreated set to 100%, as described in the Fig. 2 legend. Data shown are the mean ± SD from three independent experiments. **P < 0.01, ***P < 0.001, **** < P < 0.0001; unpaired t-test.

### Pyr01 reduces HIV-1 Env incorporation into virions in T cells

All the above experiments were performed in 293T cells. To extend our results to T cells, we next investigated whether NNRTI-induced defects in Env incorporation also occur in the SupT1 T-cell line. SupT1 cells were spinoculated with HIV-1 particles produced in 293T cells. One day postinfection, the cells were divided into three equal parts and left untreated or treated with Pyr01. Cell and viral lysates were collected 18 h later. Pyr01 treatment resulted in a substantial reduction in cellular Env expression (Fig. S9). To minimize cytotoxicity, cells were treated with Pyr01 in the presence of the caspase-1 and caspase-4 inhibitor VX-765. Consistent with our findings in 293T cells, under these conditions Pyr01 markedly reduced Env incorporation into virions (Fig. 7B) without altering cellular Env expression levels (Fig. 7A). These results demonstrate that the NNRTI-induced defect in Env incorporation is not restricted to 293T cells but also occurs in T cells.

**Fig. 7.**
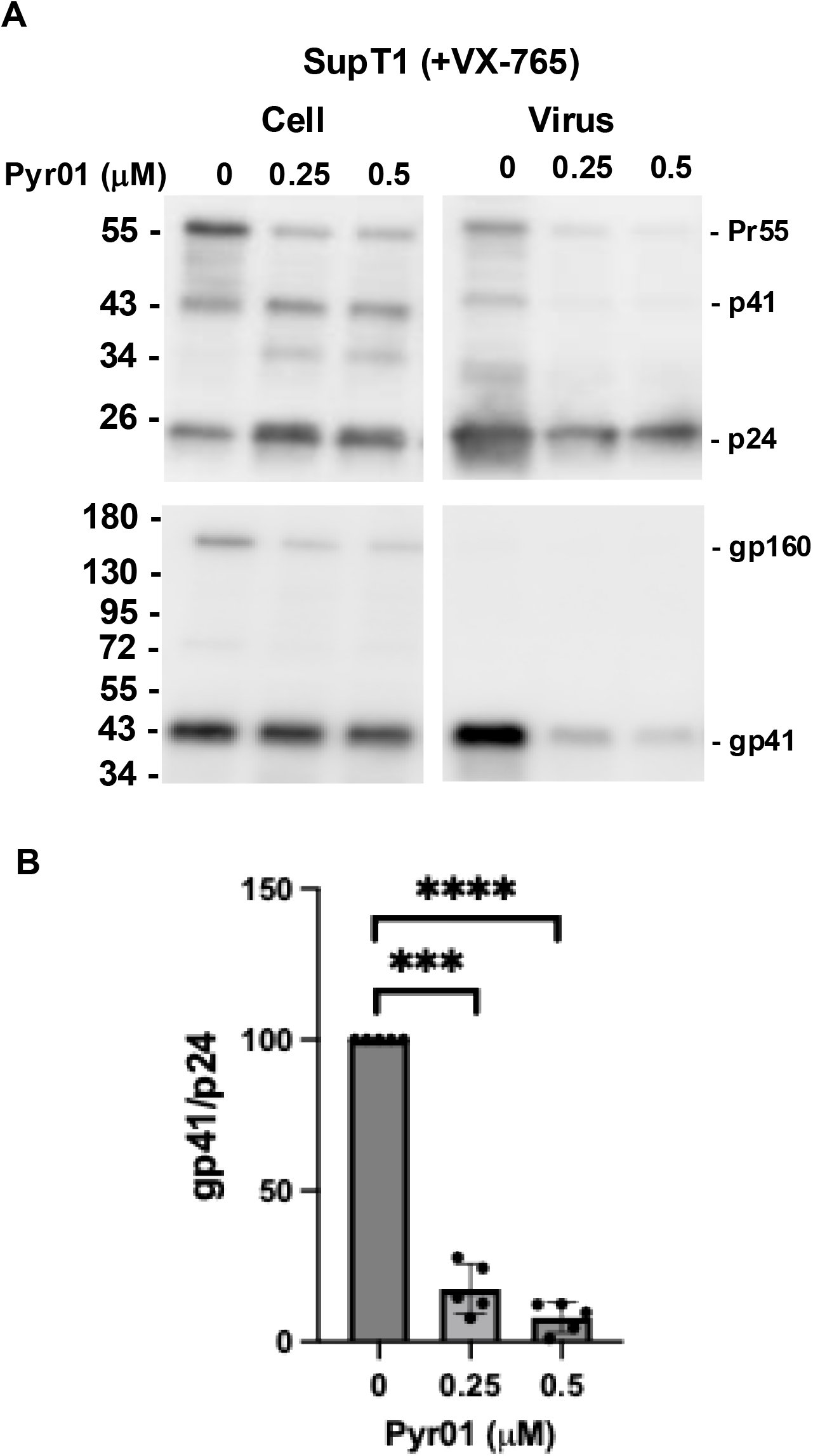
Pyr01 reduces HIV-1 Env incorporation into virions in T cells. **(A)** SupT1 cells were spinoculated with NL4-3 virus produced in 293T cells at 1,200 x g for 90 min. One day postinfection, cells were treated with the indicated concentrations of Pyr01 in the presence of caspase-1 inhibitor VX-765 (33 mg/ml). After 18 h of treatment, culture supernatants were collected, and cell and virus lysates were analyzed by western blotting using HIV-Ig and anti-gp41 2F5 antibodies, as described in the Fig. 2 legend. The positions of molecular mass standards (in kDa) are indicated on the left of each blot. **(B)** Virion-associated gp41 and p24 levels were quantified and the ratio of gp41 to p24 (CA) protein was calculated with untreated set to 100%, as described in the Fig. 2 legend. Data shown are the mean ± SD of five replicates from three independent experiments. ***P < 0.001, **** < P < 0.0001; unpaired t-test.

### Premature PR activation induced by PR-Lz and GagPol overexpression also reduces HIV-1 Env incorporation

Premature activation of HIV-1 PR can be induced by mechanisms other than NNRTI-mediated GagPol dimerization. For example, fusion of a leucine zipper (Lz) domain to the C-terminus of PR (PR-Lz) promotes GagPol dimerization and premature PR activation^60^ as does increasing the ratio of GagPol to Gag^61–63^. We therefore investigated whether premature PR activation induced by these independent approaches, like NNRTI treatment, also impairs Env incorporation into HIV-1 virions. To induce premature PR activation via PR-Lz, the pNL4-3 molecular clone was co-transfected with increasing amounts of the PR-Lz construct at NL4-3:PR-Lz plasmid ratios of 10:1, and 7.5:1. At these low PR-LZ inputs, virus particle production was only slightly reduced, but increasing PR-Lz expression resulted in a dose-dependent reduction in Env incorporation into virions (Fig. 8). Likewise, Gag and GagPol were co-expressed using NL4-3/55X and NL4-3/GagPol-FS, respectively, at NL4-3/55X:NL4-3/GagPol-FS plasmid ratios of 10:1, and 7.5:1. Increasing the ratio of GagPol to Gag also markedly reduced Env incorporation, consistent with enhanced premature PR activation (Fig. 8). Because premature PR activation also reduced cellular Env expression in these overexpression experiments, virion-associated gp41 levels were normalized to cellular Env expression. Together, these findings demonstrate that impaired Env incorporation is not unique to NNRTI treatment but is a general consequence of premature HIV-1 PR activation resulting from enhanced GagPol dimerization.

**Fig. 8.**
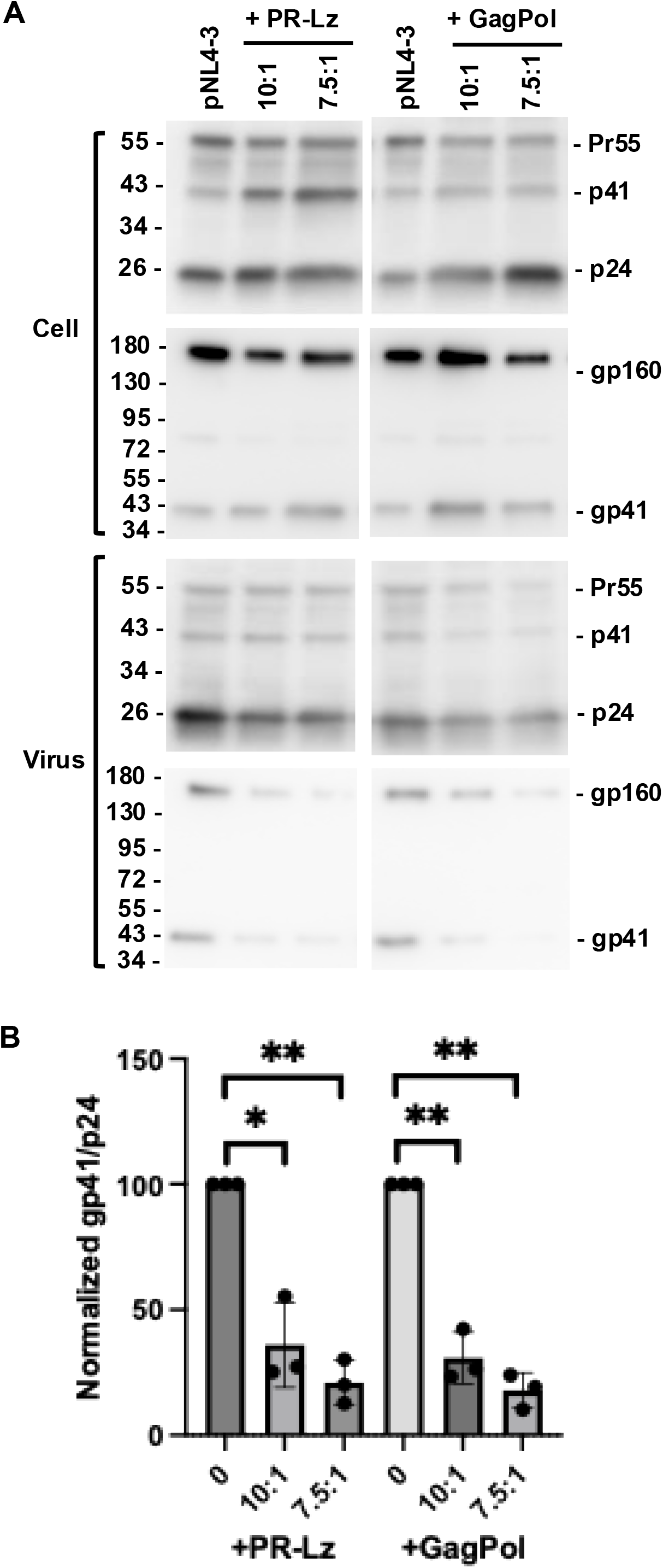
Premature PR activation induced by co-expressing PR-Lz or GagPol reduces HIV-1 Env incorporation. **(A)** 293T cells were transfected with pNL4-3 in the presence or absence of the PR-Lz construct or a GagPol expression construct at pNL4-3-to-PR-Lz/GagPol ratios of 10:1, 7.5:1 or 5:1. The medium was replaced 6-8 h posttransfection, and culture supernatants were collected 16-20 h later. Cell and virus lysates were analyzed by western blotting using HIV-Ig and anti-gp41 2F5, as described in the Fig. 2 legend. The positions of molecular mass standards (in kDa) are indicated on the left of each blot. Cell-and virion-associated gp41 levels and virion-associated p24 levels were quantified. In experiments in which PR-Lz or GagPol overexpression reduced cellular gp41 levels, virion-associated gp41 was normalized first to cellular gp41 and then to virion-associated p24 **(B)**. The resulting values were expressed relative to the corresponding control. Data represent the mean ± SD from three independent experiments. *P < 0.05, **P < 0.01; unpaired t-test.

### Pyr01 treatment reduces the levels of mature Pol products in HIV-1 virions

Premature activation of PR during virus assembly would be expected to result in premature cleavage of both Gag and GagPol, potentially resulting in loss of Pol products from the nascent virion. To investigate this possibility, we examined virion-associated Pol protein levels. Indeed, Pyr01 treatment markedly reduced the levels of mature Pol proteins IN, RT, and PR in virions (Fig. 9A). When normalized to virion-associated p24, IN levels were reduced by more than 10-fold, RT by approximately 3-fold, and PR by approximately 8-fold (Fig. 9B). To determine whether these reductions in virion-associated Pol products are due to impaired incorporation of the GagPol precursor, we examined the effects of Pyr01 on virion-associated GagPol under conditions in which PR activity was blocked. Virus-producing 293T cells expressing WT HIV-1 were treated with Pyr01 in the presence or absence of the PR inhibitor SQV. In parallel, cells expressing a catalytically inactive PR mutant [PR(-)] were treated with Pyr01. In the presence of either pharmacologic or genetic inhibition of PR activity, Pyr01 did not reduce virion-associated GagPol levels (Fig. S10) indicating that the reduction in the incorporation of Pol proteins caused by Pyr01 treatment is not due to a defect in the incorporation of GagPol into virions. Together, these results demonstrate that NNRTI treatment of virus-producing cells markedly reduces the levels of mature Pol proteins in HIV-1 virions, in addition to impairing Env incorporation, likely contributing to the severe loss of particle infectivity.

**Fig. 9.**
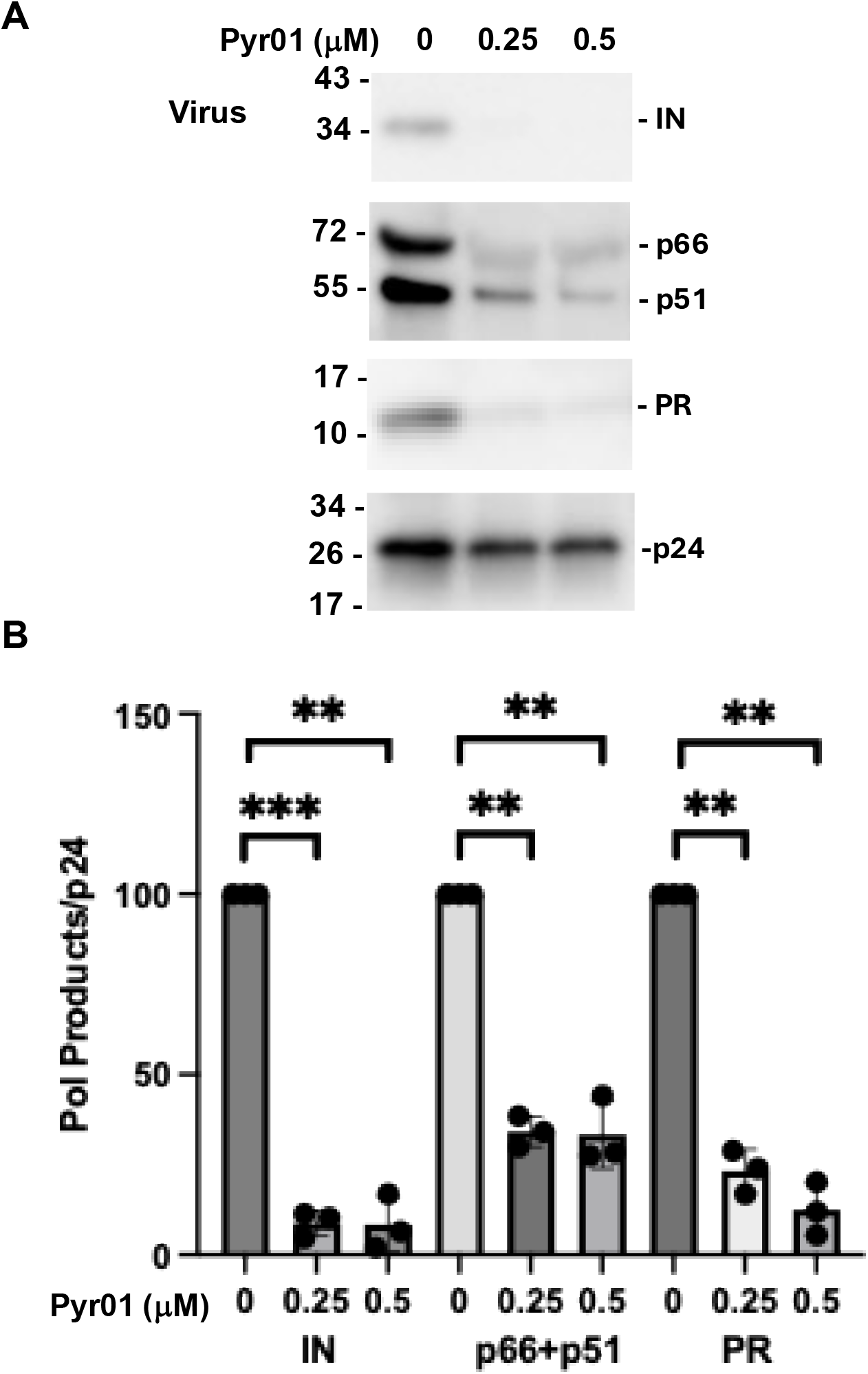
Pr01 treatment reduces the levels of mature Pol products in HIV-1 virions. **(A)** 293T cells were transfected with pNL4-3 and treated with Pyr01 at the indicated concentrations. After 16-20 h of treatment, culture supernatants were collected, and cell and virus lysates were analyzed by western blotting using antibodies against IN, RT PR, and HIV-Ig. The positions of molecular mass standards (in kDa) are indicated on the left of each blot. **(B)** Virion-associated IN, RT (p66 and p51), PR, and p24 levels were quantified. The ratios of virion-associated IN, RT, and PR to p24 were calculated with ratios from untreated samples set to 100%. Data shown are the mean ± SD from three independent experiments. **P < 0.01, ***P < 0.001; unpaired t-test.

### Premature protease activation by Pyr01 reduces viral lipid order

To determine whether Pyr01 alters viral membrane lipid packing, we labeled the virions with the solvatochromic dye, laurdan, and performed single-virus imaging of surface-immobilized particles^64,65^. We observed that mock-treated particles exhibit high lipid order [generalized polarization (GP) ∼0.32-0.4], consistent with previous measurements^26,66^. To test the sensitivity of our single-virus imaging assay, we chemically manipulated the viral membrane by depleting cholesterol with methyl-β-cyclodextrin (MβCD), which reduces lipid order. As expected, cholesterol depletion resulted in a reduction of GP (negative ΔGP values) (Fig. 10). Conversely, as expected, raising the cholesterol content of virions by treating with cholesterol-loaded MβCD (MβCD-chol) increased lipid order, resulting in positive ΔGP values (Fig. 10). To measure the effect of Pyr01 treatment on lipid order, laurdan staining was performed on virions produced from cells treated with a range of Pyr01 concentrations. Pyr01 treatment at all concentrations tested significantly reduced lipid order; at the highest concentration tested (0.5 µM), Pyr01 caused a reduction in GP (ΔGP ∼ −0.08) that exceeded that induced by cholesterol depletion of mock samples with MβCD (ΔGP ∼ −0.042; Fig. 10)^65^. These results indicate that premature activation of PR by Pyr01 in producer cells leads to viruses that exhibit loose lipid packing relative to mock-treated samples. Because the Pyr01-induced defect in Env incorporation was reversed by PR inhibition (Fig. 3), we next examined the lipid order of virions produced in the presence of Pyr01 and SQV. Importantly, Pyr01 caused only a modest additional reduction in lipid order in immature, SQV-treated viruses (ΔGP ∼ −0.021), in stark contrast to the marked decrease observed in mature, SQV-untreated particles (ΔGP ∼ −0.081). Taken together, these results indicate that Pyr01 preferentially reduces the lipid order of the HIV-1 membrane in mature (PR+) particles, supporting a role of premature PR activation in disrupting viral membrane lipid organization.

**Fig 10.**
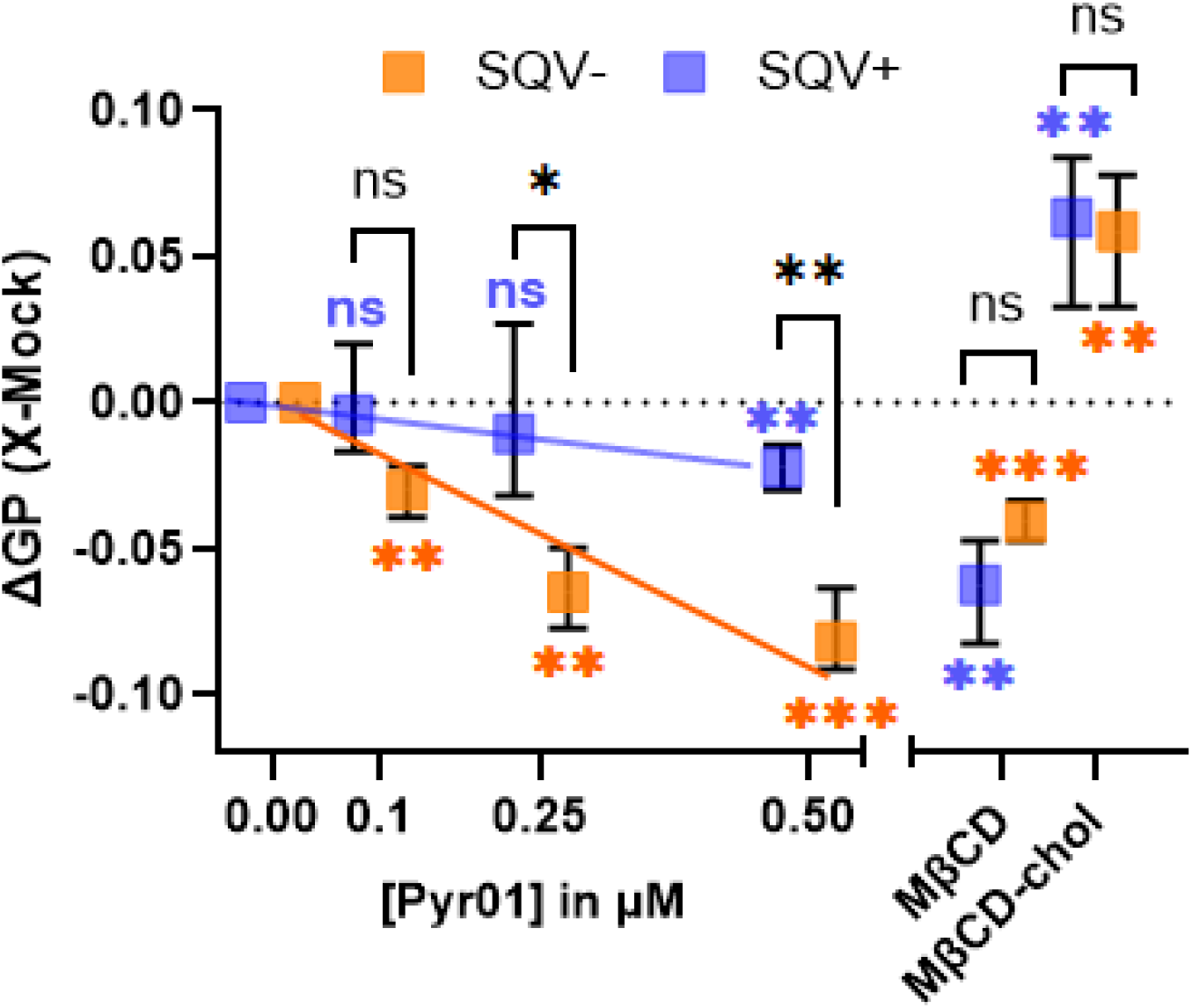
Premature PR activation induced by Pyr01 treatment reduces lipid order of the viral membrane. Quantification of changes in viral lipid order due to different treatment conditions were plotted using the ΔGP parameter. A strong reduction in lipid order upon Pyr01 treatment in saquinavir (-) (SQV-) samples (orange), but not in SQV+ (blue) samples imply selective reduction of lipid order in mature HIV-1 membranes due to premature PR activation. Chemical manipulation of the viral membrane by MβCD and MβCD-chol leads to comparable changes in lipid order in both mock-treated SQV− and 0.5µM Pyr01-treated SQV+ samples, validating the assay’s dynamic range. ΔGP for each independent panel was calculated by subtracting the median GP of mock-treated viruses from median GP of X-treated viruses (N > 100) from single-virus imaging (X ∼ 0.1-0.5 µM Pyr01, MβCD or MβCD-chol). Symbol on the graph represents mean ΔGP obtained from 3 independent replicates and error bars represent the range. Color-coded significance markers indicate differences between X and mock/control for the respective condition, whereas black significance markers denote differences between the different maturation states of viruses tested (SQV-vs SQV+). ns, not significant; p > 0.05, *; 0.05 > p > 0.01, **; 0.01 > p > 0.001; ***, p < 0.001. Statistical analysis was performed using unpaired t-test.

## Discussion

In this study, we identify a previously unrecognized effect of NNRTIs on HIV-1 virion composition and infectivity. NNRTI treatment severely reduces the incorporation of HIV-1 Env into virions, and similarly impairs the incorporation of CT-truncated HIV-1 Env and foreign viral glycoproteins, including VSV-G, SARS-CoV-2 spike and MLV Env. This defect depends on PR-mediated Gag processing, as pharmacologic inhibition or genetic inactivation of PR or mutation of Gag cleavage sites rescues Env incorporation. These findings indicate that premature PR-mediated Gag cleavage, rather than PR activity *per se*, plays a critical role in disrupting Env incorporation. Reduced Env incorporation occurs across multiple HIV-1 strains and producer cell types. Premature PR activation through alternative approaches, such as GagPol overexpression or co-expression of PR-Lz, also reduce HIV-1 Env incorporation into virions. Furthermore, Pyr01 decreases membrane lipid order in mature virions. In addition to blocking viral glycoprotein incorporation, premature PR activation also results in a substantial depletion of the Pol-derived products from virions. Together, these findings indicate that premature PR activation during virus assembly reduces HIV-1 infectivity through multiple mechanisms: impaired glycoprotein incorporation and reduced retention of Pol-derived products in nascent virions.

The requirement for PR-mediated Gag cleavage suggests that NNRTI-induced premature Gag processing, even at a limited number of cleavage sites, is central to the defect in viral glycoprotein incorporation. The immature Gag lattice provides the structural framework underlying the viral membrane^67,68^, and the MA organization within this lattice promotes Env incorporation and retention through a functional interaction with the gp41 CT ^38,39,46,69,70^. Thus, NNRTI-induced premature PR-mediated Gag processing could disrupt the MA organization within the immature Gag lattice before particle assembly is complete, thereby interfering with Env recruitment or retention at the assembly sites. However, disruption of the canonical MA-gp41 CT interaction alone is unlikely to explain the phenotype observed here, as Pyr01 also reduces the incorporation of CT-truncated HIV-1 Env, which lacks the domain that interacts with MA^38,39,41,42,46,70,71^, as well as heterologous glycoproteins, including VSV-G, SARS-CoV-2 spike and MLV Env. Moreover, the defect in glycoprotein incorporation could not be explained simply by reduced cell-surface expression as Pyr01 did not cause a corresponding decrease in the amount of HIV-1 Env or VSV-G on the plasma membrane. Together, these observations suggest a broader role for the intact immature Gag lattice in establishing and maintaining an assembly-site membrane environment that facilitates the incorporation and/or retention of diverse viral glycoproteins.

HIV-1 particles are enriched in cholesterol, sphingomyelin, and other raft-associated lipids relative to the producer-cell plasma membrane^22–24,26,27^. Early studies demonstrated that membrane-bound, multimerized Gag preferentially associates with cholesterol-dependent, detergent-resistant, raft-like membrane domains^13,14,72^, while subsequent studies showed that Gag assembly actively reorganizes the lipid and protein environment at assembly sites^29–31,73,74^.

Premature PR-mediated Gag processing could therefore perturb this specialized membrane organization before particle assembly is complete, potentially altering the distribution and organization of lipids and proteins at budding sites. Consistent with this model, we found that Pyr01 reduces the membrane lipid order in mature virions, indicating that premature activation of PR alters the physical organization of the viral membrane. Such changes in the properties of the viral membrane could provide a common mechanism for the reduced incorporation of structurally diverse viral envelope glycoproteins observed in our study. Notably, Pyr01 reduces the incorporation of glycoproteins with different membrane-domain preferences, including raft-associated full-length HIV-1 Env^13,75,76^ and MLV Env^77,78^, and non-raft-associated CT-truncated HIV-1 Env^79^, and VSV-G^78^. It also reduces the incorporation of SARS-CoV-2 spike, whose membrane organization depends on cholesterol^80^. These findings suggest that impaired glycoprotein incorporation is not restricted to proteins reportedly associated with lipid rafts. Although the precise lipid changes responsible for this phenotype remain to be defined, our findings support a model in which premature Gag processing disrupts the membrane environment established during HIV-1 assembly, thereby creating conditions less favorable for glycoprotein recruitment and/or retention.

Another consequence of premature PR activation is the depletion of Pol-derived products from virions, which may result from premature GagPol processing before particle release is complete. HIV-1 PR activation is normally coordinated with virus assembly and budding, with studies demonstrating that PR becomes active during the late stages of assembly, prior to particle release^11,12^. Premature proteolytic processing can impair particle production, maturation, and infectivity, highlighting the importance of precise temporal regulation of PR activity^12^; for review, see^10^. Consistent with this, delaying HIV-1 budding by disrupting p6 late-domain-mediated ESCRT recruitment allows PR activation and GagPol processing to occur before membrane scission is complete. Under these conditions, Pol-derived products can be lost from nascent particles, resulting in released virions depleted of viral enzymes^81–83^. The reduced levels of IN, RT and PR from virions produced in the presence of Pyr01 may reflect a similar mechanism. NNRTI-induced GagPol dimerization and premature PR activation could promote GagPol processing before membrane scission is complete, potentially allowing newly generated Pol products to escape from nascent particles rather than being retained in the virions. In addition to reduced envelope glycoprotein incorporation, this loss of essential viral enzymes would be expected to contribute substantially to the profound reduction in HIV-1 infectivity observed following Pyr01 treatment.

The effect of NNRTIs on Env incorporation is unlikely to be a secondary consequence of CARD8 inflammasome activation or pyroptotic cell death. Although NNRTI-induced premature HIV-1 PR activation can trigger CARD8-dependent pyroptosis^54^, impaired Env incorporation was readily observed in 293T cells, in which this pathway is not active^84^, indicating that CARD8-mediated cell death is not required for the phenotype. Moreover, Pyr01 treatment did not impair Env incorporation in the MA/p6 mutant, in which all PR cleavage sites have been mutated. Because the Pol sequence of MA/p6 is unchanged relative to WT, it is likely that Pyr01 induces premature PR activation in this mutant, suggesting that premature PR activation alone is insufficient to produce the Env incorporation defect. Rather, the uncoupling of PR activation from abrogated Env incorporation points to a specific Gag-dependent step downstream of premature PR activation. Together, these findings support a model in which NNRTI-induced GagPol dimerization and premature PR activation impair Env incorporation by perturbing the Gag lattice during HIV-1 assembly and maturation.

In conclusion, our findings support a model in which NNRTI-induced premature PR activation disrupts HIV-1 infectivity via two mechanisms. First, premature Gag processing disrupts the immature Gag lattice and perturbs the membrane environment established at the assembly site, thereby impairing the recruitment and incorporation of viral glycoproteins. Second, premature GagPol processing may generate Pol-derived products before membrane scission is complete, resulting in their inefficient retention within nascent particles and the production of virions depleted of essential viral enzymes. Together, these effects profoundly alter the membrane organization and protein composition of HIV-1 particles, providing a mechanistic explanation for their loss of viral infectivity. In summary, this study reveals previously unrecognized late-stage consequences of NNRTI-induced premature PR activation.

## Methods

### Plasmids

The full-length infectious HIV-1 molecular clone pNL4-3^85^, the Env-defective molecular clone pNL4-3/KFS^86^, the PR-inactive mutant pNL4-3/PR(-), which carries the D25N active-site mutation in HIV-1 PR^57^, the Pol deletion mutant pNL4-3delPol, which contains a 404-bp deletion within Pol generated by Tth111I/AgeI digestion^58^, the gp41 cytoplasmic-tail truncation mutant pNL4-3/CTdel-144^41^, the Vpu-defective pNL4-3delVpu^87^, and the Nef-defective pNL4-3/delNef^88^ have been described previously. The Gag cleavage-site mutants pNL-MA/CA, pNL-MA/SP1, pNL-MA/NC, and pNL-MA/p6.^59^ were kindly provided by C. Aiken (Vanderbilt University School of Medicine, Nashville, TN). The macrophage-tropic molecular clone pNL(AD8)^69^, the HIV-1 subtype C transmitted/founder (T/F) molecular clone K3016 (also known as CH185) and the subtype C molecular clone ZM247^89,90^ have also been described preciously. The VSV-G expression vector pHCMV-G has been described previously^91,92^. The amphotropic MLV Env expression vector pSVAMLVenv was obtained from N. Landau and D. Littman through the NIH HIV Reagent Program. The pcDNA3.1-SARS-CoV-2-S expression vector encoding the SARS-CoV-2 Wuhan strain spike glycoprotein was a gift from T. Gallagher (Loyola University, Chicago, IL). The PR-Lz construct^60^, in which a heterologous leucine zipper was fused to the C-terminus of HIV-1 PR, was obtained from C-T. Wang (National Yang-Ming University School of Medicine, Taipei, Taiwan). The Gag expression plasmid pNL4-3/55X was described previously^93^. To express GagPol independently of Gag, the native (-1) ribosomal frameshift site was modified to place *gag* and *pol* in the same reading frame, generating pNL4-3/GagPol-FS. The plasmid encoding HXB2 Env, the PR9ΔEnv HIV-1 based packaging vector, and the mRFP-Vpr plasmid, have been described previously^65^.

### Antibodies and chemicals

The following antibodies were obtained from the NIH HIV Reagent Program: anti-HIV-1 immunoglobulin (HIV-Ig), antibodies against HIV-1 IN, RT, PR, anti-gp41 monoclonal antibodies 2F5 and 10E8, and anti-gp120 monoclonal antibodies VRC01, 2G12, and PGT121. Antibodies against VSV-G were purchased from abcam, and the SARS-CoV-2 Spike antibody was purchased from Sino Biological. Anti-FLAG and anti-HA antibodies were purchased from Sigma-Aldrich. Anti-MLV-gp70 antibody was kindly provided by R. Gorelick (AIDS and Cancer Virus Program, Frederick National Laboratory for Cancer Research). The NNRTI Pyr01 was kindly provided by E. Arnold (Center for Advanced Biotechnology and Medicine, Rutgers Health, Piscataway, NJ) and was also purchased from MedChemExpress LLC. The Caspase-1/4 inhibitor VX-765 was purchased from Sigma-Aldrich Inc. The NNRTI EFV and the HIV-1 protease inhibitor SQV were obtained from the NIH HIV Reagent Program. Laurdan was obtained from Anaspec (Fremont, CA, USA). Anti-tubulin antibody was purchased from Sigma-Aldrich.

### Cell culture and transfection

HEK293T cells were obtained from the American Type Culture Collection (ATCC) (Manassas, VA), and TZM-bl cells were obtained from J. C. Kappes through the NIH HIV Reagent Program). These cell lines were maintained at 37 °C and 5% CO_2_ in Dulbecco’s modified Eagle’s medium (DMEM) supplemented with 10% fetal bovine serum (FBS; HyClone), 2 mM glutamine, and 1% penicillin-streptomycin. SupT1cells were cultured in Roswell Park Memorial Institute (RPMI) medium with 10% FBS, 2 mM glutamine, and 1% penicillin-streptomycin under the same conditions. For transfections, one day after plating, 293T cells were transfected with the indicated plasmid DNA using Lipofectamine 2000 (Invitrogen) according to the manufacturer’s instructions. SupT1 cells were infected via spinoculation with NL4-3 virus produced in 293T cells at 1,200 x g for 90 min. Virus-containing supernatants from HEK293T and SupT1 cells were harvested at 16-20 h after change of medium and filtered through a 0.45-μm membrane filter. An aliquot was stored for infectivity assay, and the remainder was pelleted by ultracentrifugation. Cell and virion pellets were solubilized in lysis buffer containing 10 mM iodoacetamide (Sigma-Aldrich), complete protease inhibitor tablets (Roche), 300 mM sodium chloride, 50 mM Tris-HCl (pH 7.5), and 0.5% Triton X-100 (Sigma-Aldrich).

### Western blotting

Cell and virus lysates were mixed with 6x sodium dodecyl sulfate-polyacrylamide gel electrophoresis (SDS-PAGE) sample loading buffer [600 mM Tris-HCl (pH 6.8), 30% glycerol, 12% SDS, 20 mM dithiothreitol, and 0.03% bromophenol blue] and denatured at 95 °C for 5 min. Proteins were resolved on 4–15% Tris-glycine polyacrylamide gels and transferred to PVDF membranes using the Bio-Rad Trans-Blot Turbo Transfer System according to the manufacturer’s instructions. Membranes were probed with the indicated primary antibodies followed by appropriate secondary antibodies. Protein bands were detected by chemiluminescence, imaged using a Gel Doc XR+ system (Bio-Rad), and quantified using Image Lab software version 6.0.1 (Bio-Rad) or AzureSpot (Azure Biosystems).

### Single-cycle infectivity assays

TZM-bl cells, a HeLa cell-derived reporter cell line stably expressing luciferase and β-galactosidase under the control of the HIV-1 long terminal repeat (LTR) promotor^94^, were infected with serial dilutions of virus stocks in the presence of 20 μg/mL DEAE-dextran. At 48 h post-infection, cells were lysed with Britelite luciferase reagent (PerkinElmer), and luciferase activity was measured using the GloMax Navigator microplate luminometer (Promega).

### Flow cytometry

Transfected HEK293T cells (5 X 10^5^) were resuspended in 500 µL of cold PBS. To detect cell-surface HIV-1 Env, the anti-gp120 monoclonal antibodies VRC01, 2G12, and PGT121 were pooled, each at a 1:1,000 dilution, in PBS containing 1% BSA. Similarly, an anti-VSV-G antibody was diluted 1:1,000 in PBS containing 1% BSA for detection of cell-surface VSV-G. Cells were incubated with the respective primary antibodies for 30 min and then washed twice with PBS containing 1% BSA. Cells were subsequently incubated with the appropriate Alexa Fluor 595-conjugated secondary antibody at a 1:1,000 dilution for 30 min and washed twice with PBS containing 1% BSA. Cells were then fixed in 150 µL of Cytofix/Cytoperm solution (BD Biosciences, catalog no. 554714) for 30 min and washed with 1x Perm/Wash buffer (BD Biosciences). Finally, cells were resuspended in 200 µL of PBS and analyzed using a BD LSRFortessa flow cytometer (BD Biosciences). Data were exported as FCS 3.0 files and analyzed using FlowJo version 10.9.0 (Tree Star, Inc.). A consistent gate was applied across all samples to quantify the percentage of cells expressing HIV-1 Env or VSV-G on the cell surface and to determine the mean fluorescence intensity (MFI).

### Fluorescent HIV-1 production and characterization

For laurdan staining experiments, fluorescent HIV-1 pseudoviruses containing an RFP-Vpr fiducial-marker were prepared by transfecting HEK293T cells in 6-well plates with 0.67 μg of HXB2 Env expression vector, 0.89 μg of PR9ΔEnv vector, 0.15 μg of mRFP-Vpr plasmid using JetPRIME (Polyplus-Sartorius).

After 8 h, the medium was replaced with medium containing the indicated concentrations of Pyr01 in the presence or absence of 0.5 µM SQV. Approximately 48 h post-transfection, the supernatants were collected, passed through 0.45 µm PES filters, and subsequently concentrated 10-fold using Lenti-X concentrator (Takara Bio, San Jose, CA, USA). The viruses were then aliquoted and stored at −80 °C. The p24 content of each viral preparation was determined by ELISA, as previously described^95^. Pseudovirus infectivity was measured using TZM-bl cells as described above. To gauge PR activity in individual pseudoviral preparations, the samples were analyzed using western blotting as described above.

### Single-virus lipid order measurements

Pseudovirus stocks (∼0.2-0.5 ng of p24) were thawed and stained with freshly prepared laurdan at final concentration of 28 µM. This mixture was then incubated at 37 °C for 10 min, diluted 10-fold, and attached to a poly-D-lysine (PDL)-coated coverglass (Cellvis, Mountain View, CA, USA) for single-virus imaging. For cholesterol depletion or supplementation, the immobilized viruses were first rinsed 3 times with PBS and then incubated with 0.15 mM methyl-β-cyclodextrin (MβCD) or 0.25 mM MβCD-cholesterol (MβCD-chol) complex, respectively, for 30 minutes at 37 °C^65^. The viruses were then rinsed at least 3 times with PBS prior to imaging. Fluorescent viruses were imaged using a Zeiss LSM880 microscope (Carl Zeiss Microimaging, Jena, Germany) as previously described^65^. Briefly, mRFP puncta were used to extract precise coordinate information of immobilized viruses, and laurdan signal was measured exclusively from these puncta through wavelet-based localization algorithms via ICY software^65^. This enabled the removal of non-specific laurdan signal. The lipid order of viral particles was computed using the generalized polarization parameter (GP) of laurdan using the following formula, where I443 nm and I513 nm are laurdan fluorescence emission signals at respective wavelengths, when excited with 405 nm light.

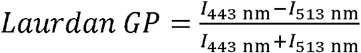

## Acknowledgments

We thank members of the Freed lab for helpful discussion and critical review of the manuscript. Work in the Freed lab is supported by the Intramural Research Program of the Center for Cancer Research, National Cancer Institute, NIH. Work in the Melikyan lab was supported by the NIH R37 AI150453 and R01 AI189249 grants.

## Author contributions

A.A.W. and E.O.F designed the research; A.A.W., H.I, and G.R. performed the experiments; A.A.W., G.R., G.B.M. and E.O.F. analyzed the data; A.A.W. and E.O.F. wrote the manuscript. All authors read and approved the manuscript.

