## Supplemental Figures for "NON-NUCLEOSIDE REVERSE TRANSCRIPTASE INHIBITORS BLOCK THE INCORPORATION OF VIRAL GLYCOPROTEINS INTO HIV-1 VIRIONS"

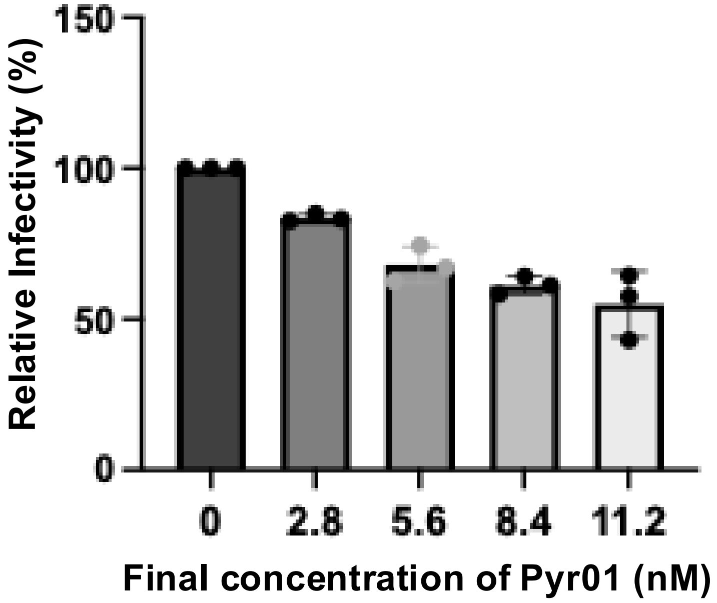

150
100
50
0
Relative Infectivity (%)
0
2.8
5.6
8.4
11.2
Final concentration of Pyr01 (nM)


Figure S1

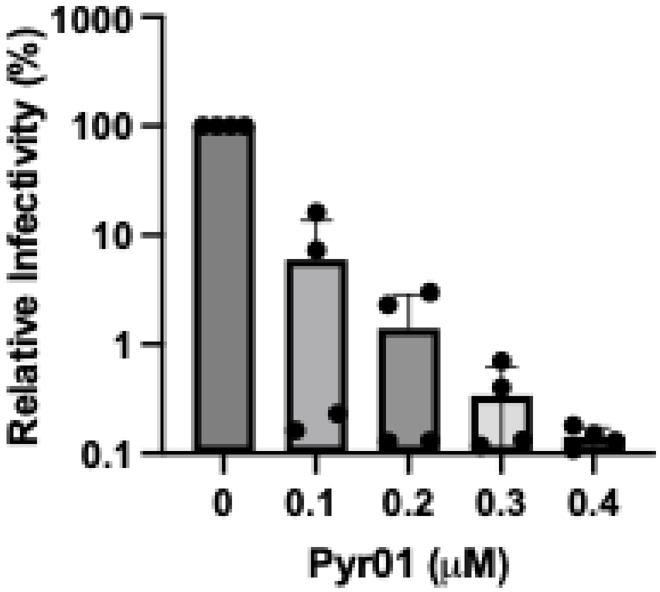


Figure S2

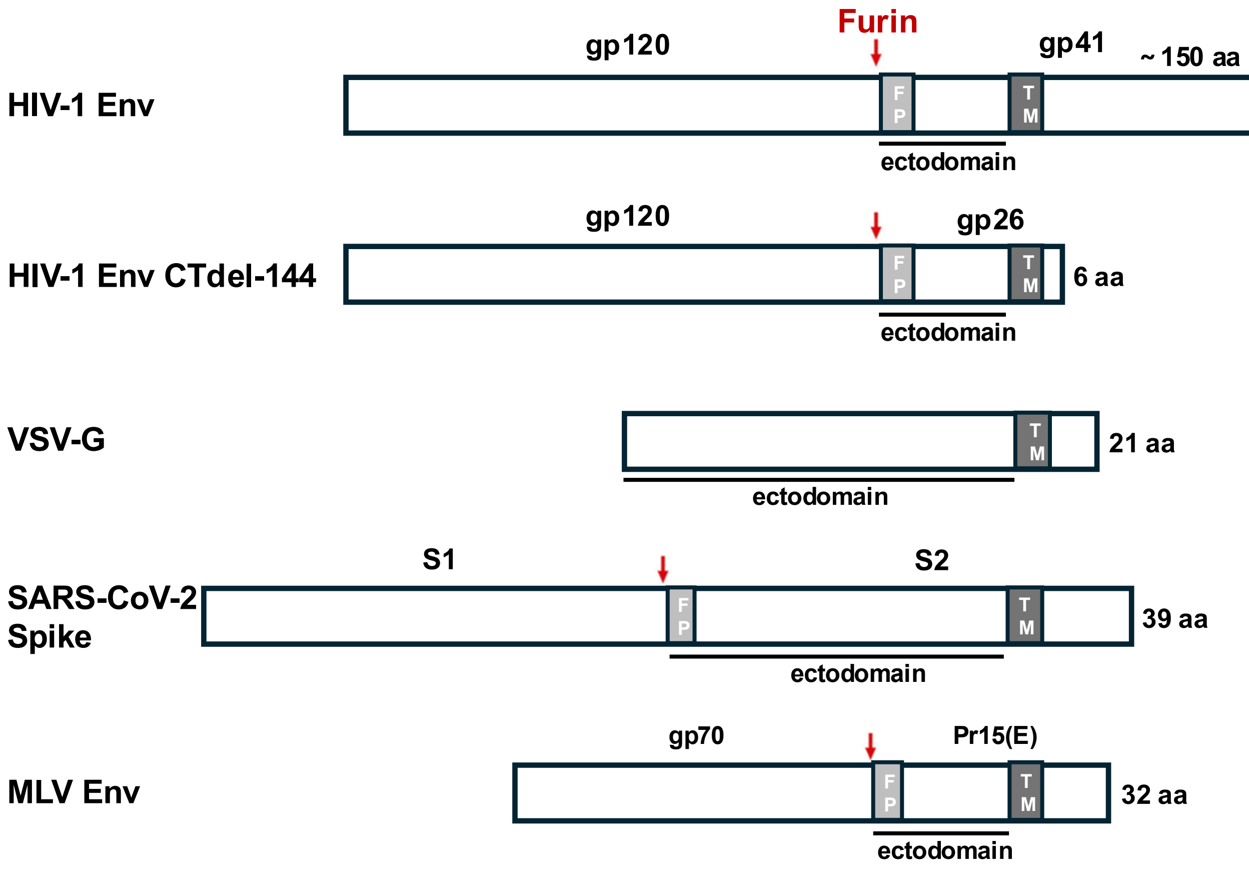
Furin
gp120
gp41
~ 150 aa
HIV-1 Env
F
P
T
M
ectodomain
gp120
gp26
HIV-1 Env CTdel-144
6 aa
ectodomain
VSV-G
21 aa
ectodomain
S1
S2
SARS-CoV-2
Spike
39 aa
ectodomain
gp70
Pr15(E)
MLV Env
32 aa
ectodomain

Figure S3

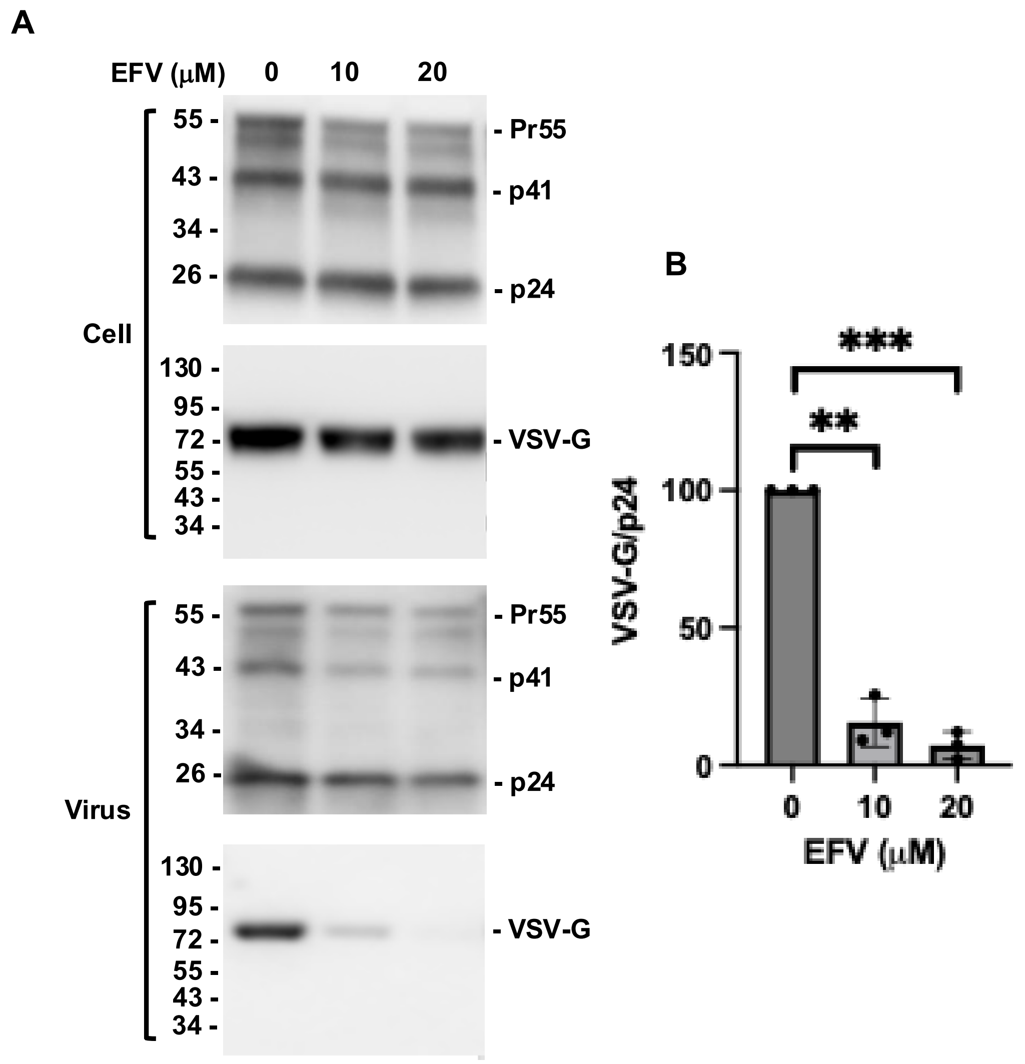
A
EFV (μM)
0
10
20
Cell
55 -
43 -
34 -
26 -
- Pr55
- p41
- p24
130 -
95 -
72 -
55 -
43 -
34 -
- VSV-G
Virus
55 -
43 -
34 -
26 -
- Pr55
- p41
- p24
130 -
95 -
72 -
55 -
43 -
34 -
- VSV-G
B
VSV-G/p24
150
100
50
0
***
**
0
10
20
EFV (μM)

Figure S4

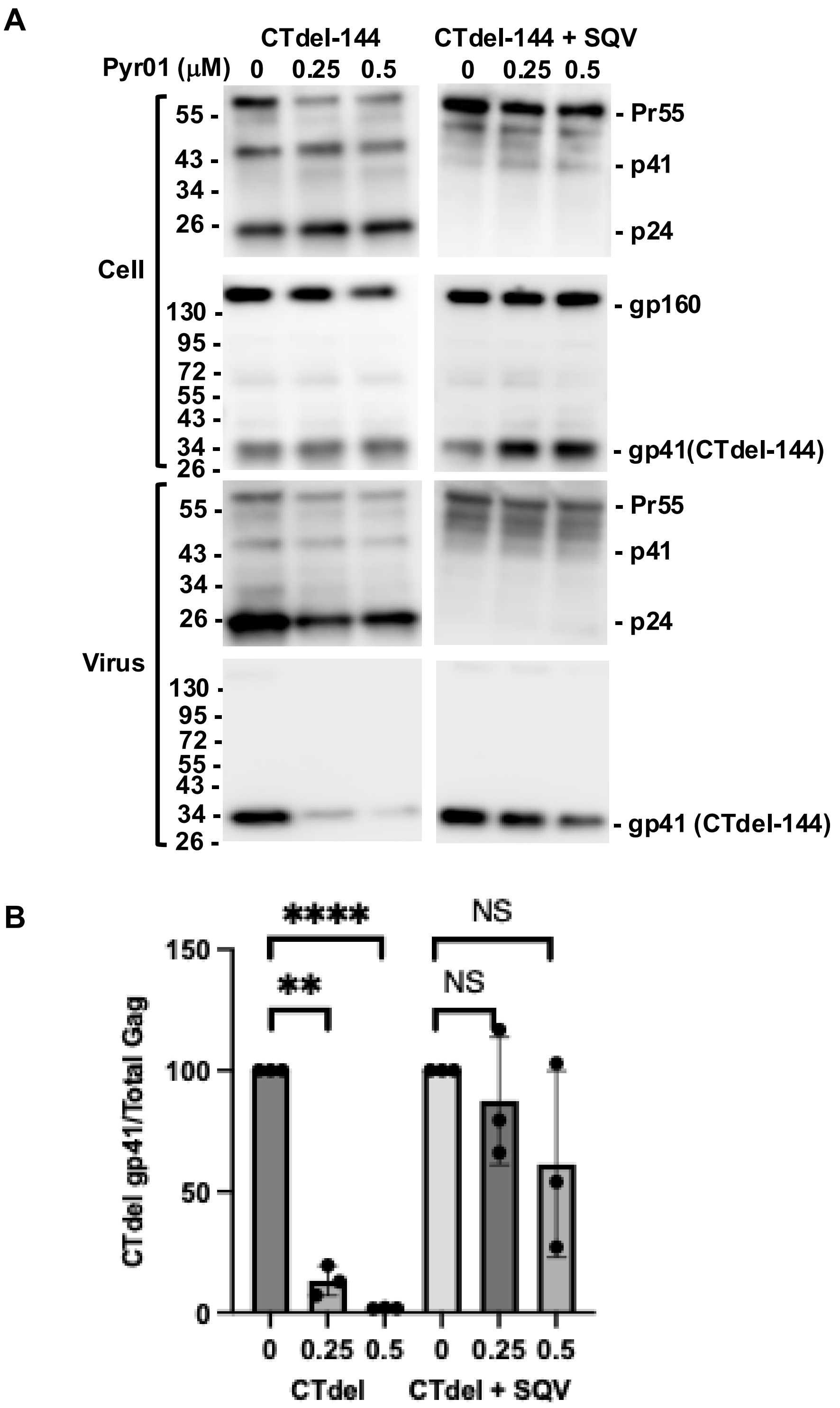
A
CTdel-144
CTdel-144 + SQV
Pyr01 (μM) 0 0.25 0.5 0 0.25 0.5
Cell
Virus
55 -
43 -
34 -
26 -
130 -
95 -
72 -
55 -
43 -
34 -
26 -
- Pr55
- p41
- p24
- gp160
- gp41(CTdel-144)
- gp41 (CTdel-144)
B
CTdel gp41/Total Gag
150
100
50
0
****
**
NS
NS
0 0.25 0.5 0 0.25 0.5
CTdel
CTdel + SQV

Figure S5

A

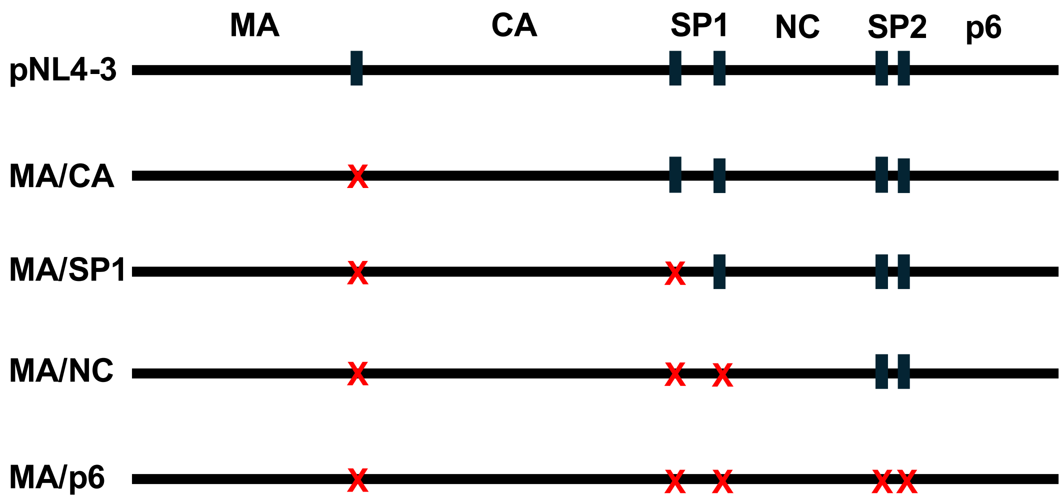
MA
CA
SP1
NC
SP2
p6
pNL4-3
MA/CA
MA/SP1
MA/NC
MA/p6

B

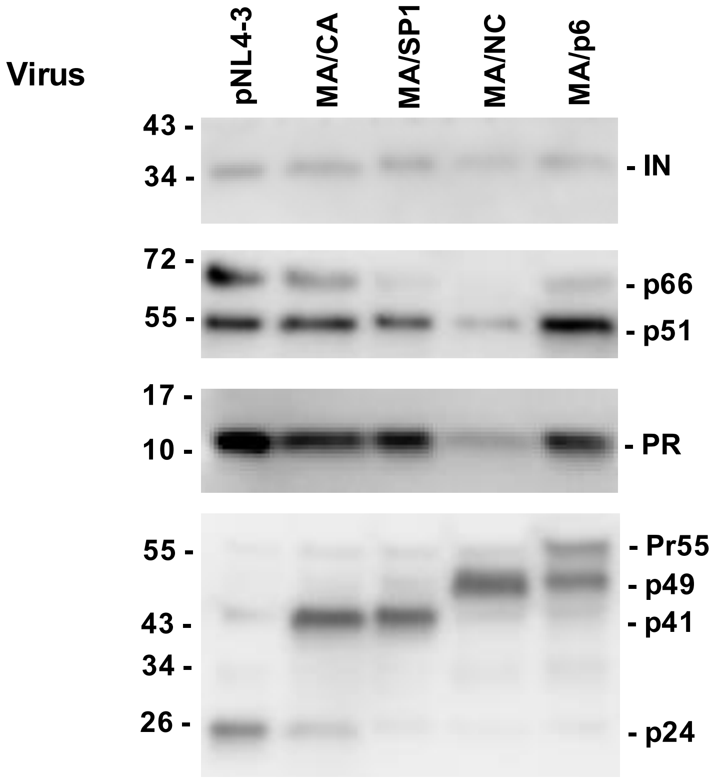
Virus
pNL4-3
MA/CA
MA/SP1
MA/NC
MA/p6
43 -
34 -
- IN
72 -
55 -
- p66
- p51
17 -
10 -
- PR
55 -
43 -
34 -
26 -
- Pr55
- p49
- p41
- p24

Figure S6

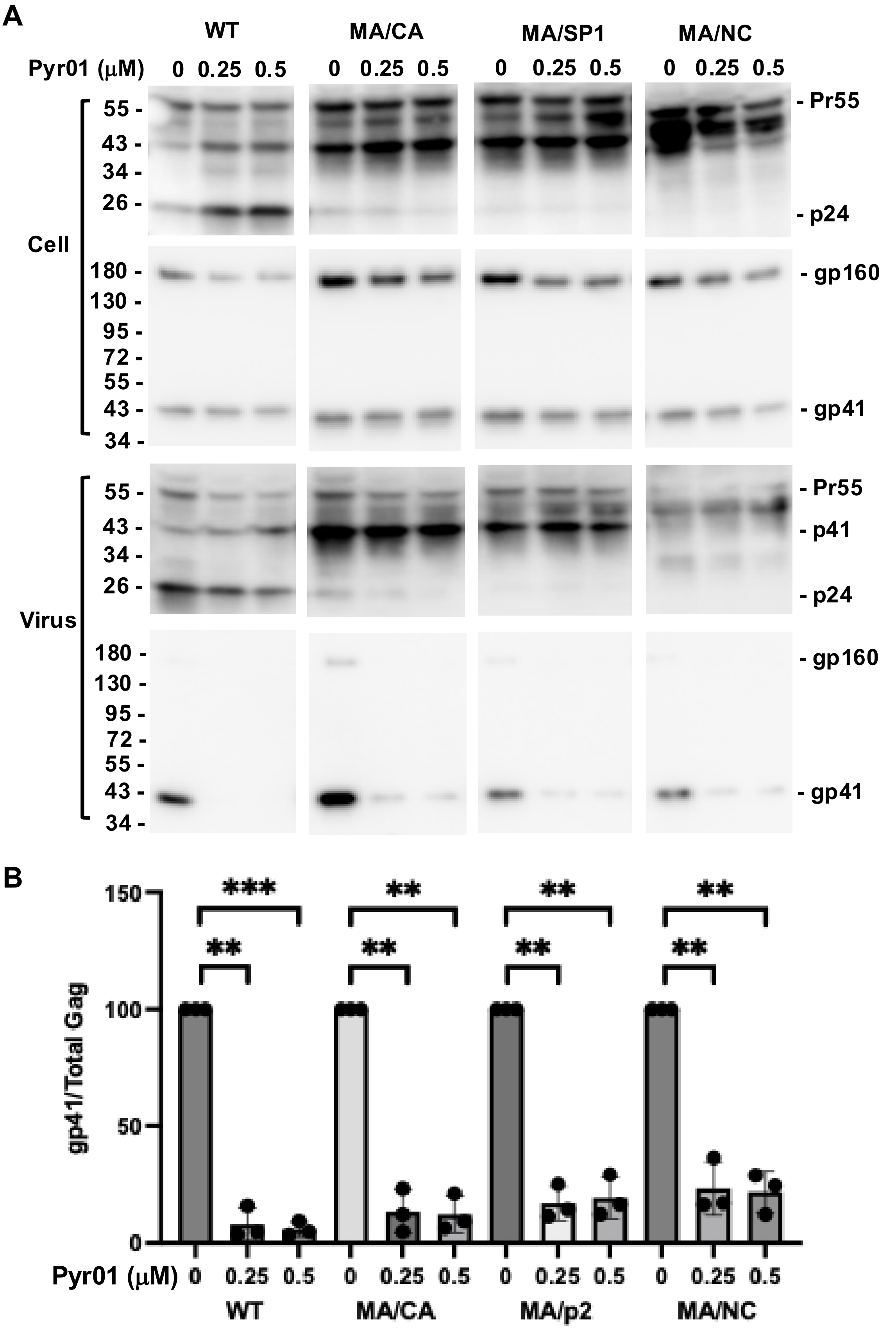
A
WT
MA/CA
MA/SP1
MA/NC
Pyr01 (μM)
0 0.25 0.5
Cell
Virus
55 -
43 -
34 -
26 -
180 -
130 -
95 -
72 -
- Pr55
- p24
- gp160
- gp41
- p41
B
gp41/Total Gag
150
100
50
0
***
**
Pyr01 (μM)
0 0.25 0.5
WT
MA/CA
MA/p2
MA/NC

Figure S7

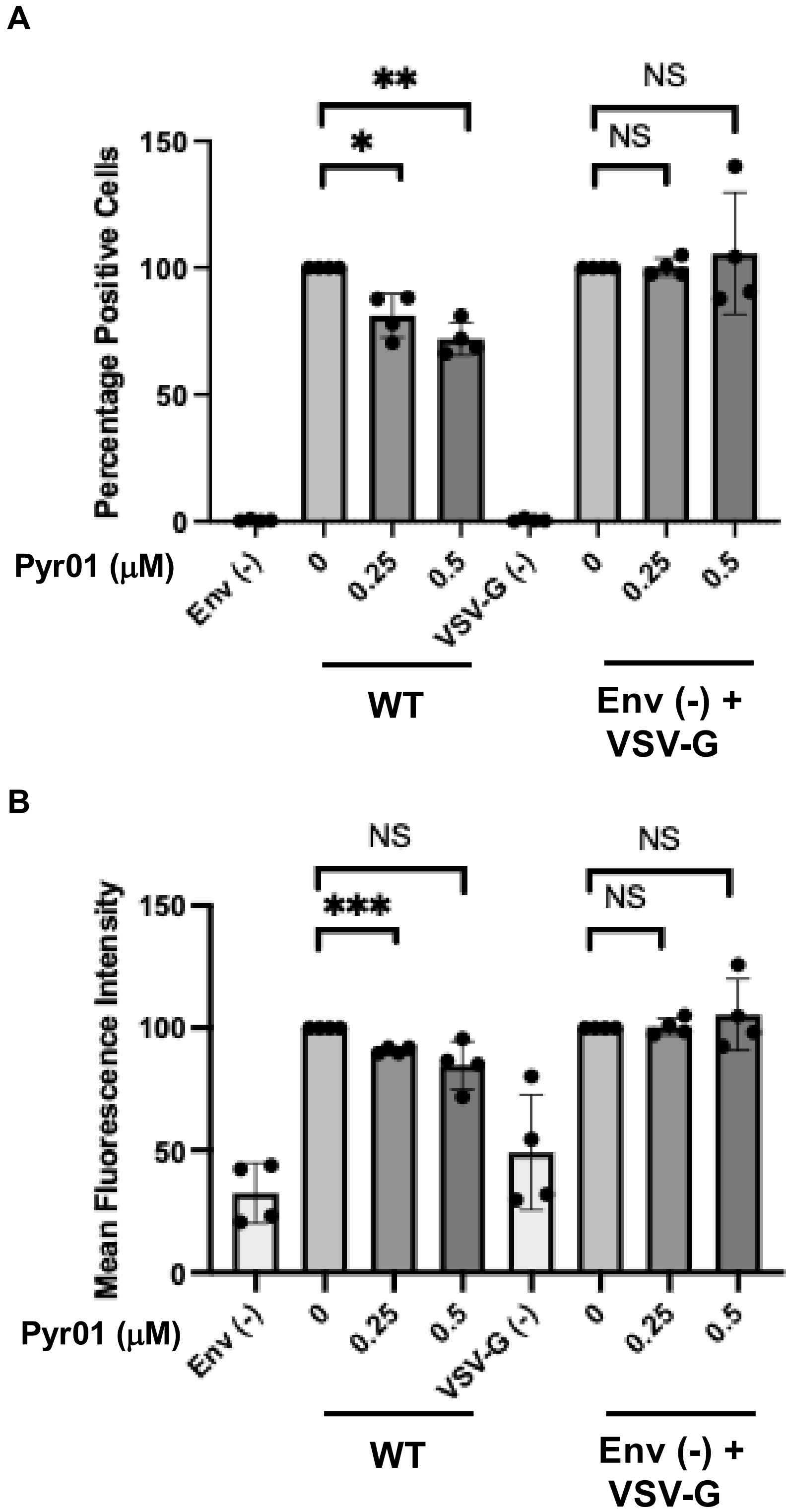
A
**
*
NS
NS
Percentage Positive Cells
150
100
50
0
Pyr01 (μM)
Env (-)
0
0.25
0.5
VSV-G (-)
0
0.25
0.5
WT
Env (-) +
VSV-G
B
NS
***
NS
NS
Mean Fluorescence Intensity
150
100
50
0
Pyr01 (μM)
Env (-)
0
0.25
0.5
VSV-G (-)
0
0.25
0.5
WT
Env (-) +
VSV-G

Figure S8

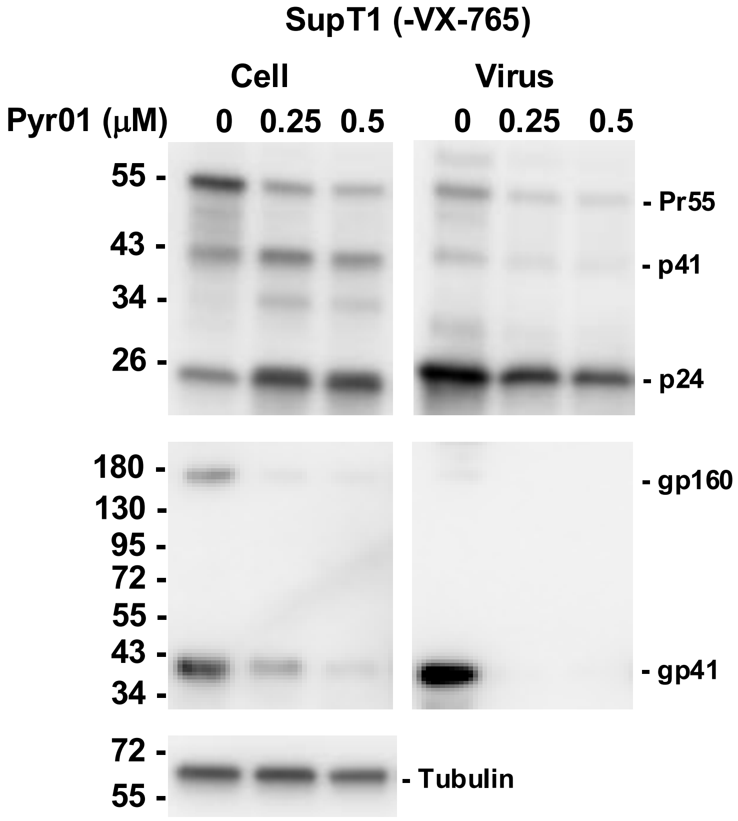
SupT1 (-VX-765)
Cell
Virus
Pyr01 (μM)
0
0.25
0.5
0
0.25
0.5
55 -
43 -
34 -
26 -
- Pr55
- p41
- p24
180 -
130 -
95 -
72 -
55 -
43 -
34 -
- gp160
- gp41
72 -
55 -
- Tubulin

Figure S9

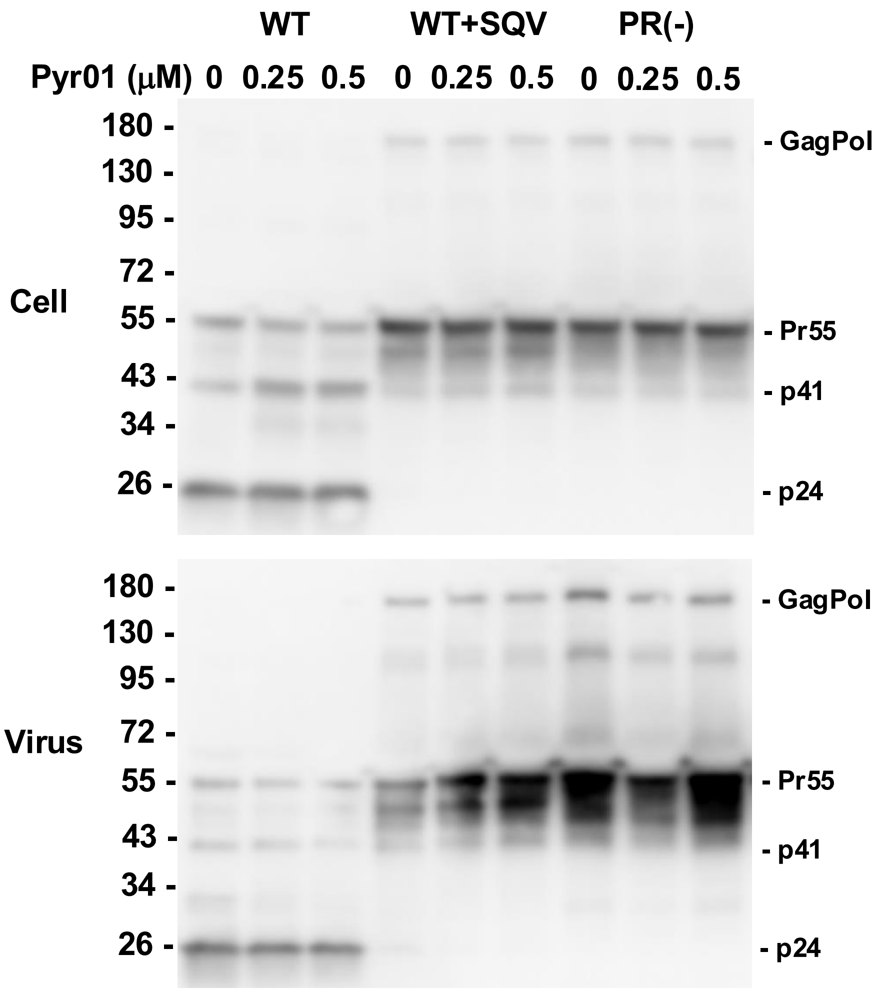
WT
WT+SQV
PR(-)
Pyr01 (μM) 0 0.25 0.5 0 0.25 0.5 0 0.25 0.5
Cell
Virus
180 -
130 -
95 -
72 -
55 -
43 -
34 -
26 -
- GagPol
- Pr55
- p41
- p24

Figure S10
