## Supplementary Figure Legends for "NON-NUCLEOSIDE REVERSE TRANSCRIPTASE INHIBITORS BLOCK THE INCORPORATION OF VIRAL GLYCOPROTEINS INTO HIV-1 VIRIONS"

and Eric O. Freed^1*^

^1^Virus-Cell Interaction Section, HIV Dynamics and Replication Program, Center for Cancer Research, National Cancer Institute, Frederick, MD 21702, USA

^2^Department of Pediatrics, Division of Infectious Diseases, Emory University School of Medicine, Atlanta, GA 30322, USA

**Supplementary Figure Legends**

**Fig. S1. Effect of Pyr01 addition during infection.** Pyr01 was added to TZM-bl cells at the indicated final concentrations, followed by infection with HIV-1 virions. Luciferase activity was measured 48 h postinfection, and infectivity in the absence of Pyr01 was set to 100%. Data represent the mean ± SD from three independent experiments.

**Fig. S2. Pyr01 treatment during virus production markedly impairs the infectivity of VSV-G pseudotyped HIV-1 virions.** 293T cells were co-transfected with the Env-defective HIV-1 molecular clone pNL4-3/KFS [Env(-)] and a VSV-G expression vector. After 6-8 h posttransfection, cells were treated with Pyr01 at 0, 0.1, 0.2, 0.3 and 0.4 μM. After 16-20 h of treatment, culture supernatants were collected and TZM-bl cells were infected with serial dilutions of the virus-containing supernatants. Luciferase activity was measured 48 h postinfection, and the infectivity of virus produced in the absence of Pyr01 was set to 100%. Data shown are the mean ± SD from four independent experiments.

**Fig. S3. Viral envelope glycoproteins used in this study.** Surface and transmembrane subunits of WT and gp41 cytoplasmic tail-truncated HIV-1 Env CTdel144 HIV-1 Env (gp120 and gp41), VSV-G, SARS-CoV-2 Spike (S1 and S2), and MLV Env [gp70 and Pr15(E)], are indicated. Furin cleavage is indicated by a red arrow; the fusion peptide (FP), transmembrane domains (TM), and ectodomains are labeled. The approximate length of the cytoplasmic tails of each envelope glycoprotein is indicated in amino acids (aa).

**Fig. S4.** **EFV reduces the incorporation of VSV-G into HIV-1 particles. (A)** 293T cells were transfected with the Env-defective molecular clone pNL4-3/KFS [Env(-)] and VSV-G expression vector. Six to eight h posttransfection, cells were treated with EFV at 0, 10 and 20 μM. After 16-20 h of treatment, culture supernatants were collected, and cell and virus lysates were analyzed by western blotting using HIV-Ig and an anti-VSV-G antibody. The positions of molecular mass standards (in kDa) are indicated on the left of each blot. **(B)** Virion-associated VSV-G and p24 levels were quantified and the ratio of VSV-G to p24 protein was calculated with untreated set to 100%, as described in the Fig. 2 legend. Data represent the mean ± SD from three independent experiments. **P < 0.01, ***P < 0.001; unpaired t-test.

**Fig. S5.** **Inhibition of PR activity with saquinavir reverses the Pyr01-mediated defect in HIV-1 CTdel-144 Env incorporation. (A)** 293T cells were transfected with the gp41 cytoplasmic-tail truncation mutant pNL4-3/CTdel-144. Six to eight h posttransfection, cells were treated with Pyr01 at 0, 0.25 and 0.5 μM in the presence or absence of the PR inhibitor saquinavir (SQV) (0.5 μM). After 16-20 h of treatment, culture supernatants were collected, and cell and virus lysates were analyzed by western blotting using HIV-Ig, and the gp41 antibody 2F5. The positions of molecular mass standards (in kDa) are indicated on the left of each blot. (**B)** Virion-associated gp41 and p24 levels were quantified and the ratio of gp41 to p24 protein was calculated with untreated set to 100%, as described in the Fig. 2 legend Data shown are the mean ± SD from three independent experiments. NS - not significant; **P < 0.01; ****P < 0.0001; unpaired t-test.

**Fig. S6. HIV-1 PR remains active in Gag cleavage-site mutants. (A)** Schematic representation of the Gag cleavage site mutants tested. Mutated cleavage sites are indicated by red Xs. **(B)** 293T cells were transfected with WT pNL4-3 or the Gag cleavage-site mutants pNL-MA/CA (MA/CA), pNL-MA/SP1 (MA/SP1), pNL-MA/NC (MA/NC), or pNL-MA/p6 (MA/p6). Six to eight h posttransfection, the medium was replaced. Virus supernatants were collected and analyzed by western blotting using antibodies against IN, RT, PR, and HIV-Ig. The positions of molecular mass standards (in kDa) are indicated on the left of each blot.

**Fig. S7.** **Partial inhibition of Gag cleavage does not rescue the Pyr01-induced defect in Env incorporation. (A)** 293T cells were transfected with WT pNL4-3 (WT) or the Gag cleavage-site mutants pNL-MA/CA (MA/CA), pNL-MA/SP1 (MA/SP1), or pNL-MA/NC (MA/NC). Six to eight h posttransfection, cells were treated with Pyr01 at the indicated concentrations. After 16-20 h of treatment, culture supernatants were collected, and cell and virus lysates were analyzed by western blotting using HIV-Ig and anti-gp41 antibody 2F5, as described in the Fig. 2 legend. The positions of molecular mass standards (in kDa) are indicated on the left of each blot. **(B)** Virion-associated gp41 and p24 levels were quantified and the ratio of gp41 to p24 protein was calculated with untreated set to 100%, as described in the Fig. 2 legend. Data shown are the mean ± SD from three independent experiments. **P < 0.01, ***P < 0.001; unpaired t-test.

**Fig. S8. Effect of Pyr01 on cell-surface expression of HIV-1 Env and VSV-G.** 293T cells were transfected with WT pNL4-3 (WT) or co-transfected with the Env-defective molecular clone pNL4-3/KFS [Env(-)] and a VSV-G expression vector. Six h posttransfection, cells were treated with Pyr01 at 0, 0.25 and 0.5 μM. After 16 h of treatment, cells were collected and incubated with a pool of the anti-gp120 monoclonal antibodies VRC01, 2G12, and PGT121 or with anti-VSV-G antibody, as described in the Methods. After washing, cells were incubated with the appropriate Alexa Fluor 595-conjugated secondary antibody, washed, fixed and analyzed by flow cytometry. The percentage of cells expressing HIV-1 Env or VSV-G on the cell surface **(A)** and mean fluorescence intensity **(B)** of cell-surface Env and VSV-G were determined using FlowJo version 10.9.0 software. Values for untreated cells were set to 100%. Data shown are the mean ± SD from four independent experiments. NS - not significant; *P < 0.05; **P < 0.01; ***P < 0.001; unpaired t-test.

**Fig. S9.** **Pyr01 reduces HIV-1 Env expression in SupT1 cells in the absence of a capase-1 inhibitor.** SupT1 cells were spinoculated with NL4-3 virus produced in 293T cells at 1,200 x g for 90 min. One day postinfection, cells were treated with the indicated concentrations of Pyr01 (in the absence of caspase-1 inhibitor VX-765). After 18 h of treatment, culture supernatants were collected, and cell and virus lysates were analyzed by western blotting using HIV-Ig and anti-gp41 2F5 antibodies, as described in the Fig. 2 legend. Cell lysates were also probed with an anti-tubulin antibody as a loading control. The positions of molecular mass standards (in kDa) are indicated on the left of each blot. Data shown are representative of three independent experiments.

**Fig. S10**. **Pyr01 treatment does not reduce GagPol incorporation into virions.** (A) 293T cells were transfected with WT pNL4-3 (WT), or a PR-inactive mutant [PR(-)]. Six to eight h posttransfection, cells were treated with 0, 0.25 and 0.5 μM Pyr01 in the presence or absence of 0.5 μM of the protease inhibitor SQV. After 16-20 h of treatment, culture supernatants were collected and cell and virus lysates were analyzed by western blotting using HIV-Ig, as described in the Fig. 2 legend. The positions of molecular mass standards (in kDa) are indicated on the left of each blot. Data shown are representative of three independent experiments.
